# A Tough Biointerface in Human Knee Empowered by Dynamic Phase-transforming Minerals in Collagenous Matrix

**DOI:** 10.1101/2024.08.03.606023

**Authors:** Wenyue Li, Xiaozhao Wang, Renwei Mao, Dong Li, Hongxu Meng, Ru Zhang, Jinghua Fang, Zhengzhong Kang, Boxuan Wu, Weiwei Ma, Xudong Yao, Chang Xie, Rui Li, Jin Wang, Xiao Chen, Xihao Pan, Weiqiu Chen, Wangping Duan, Huajian Gao, Hongwei Ouyang

**Affiliations:** Department of Sports Medicine of the Second Affiliated Hospital, and Liangzhu Laboratory, Zhejiang University School of Medicine; Hangzhou, China; Dr. Li Dak Sum & Yip Yio Chin Center for Stem Cells and Regenerative Medicine, Zhejiang University School of Medicine; Hangzhou, China; Zhejiang University-University of Edinburgh Institute, Zhejiang University School of Medicine; Haining, China; China Orthopedic Regenerative Medicine Group (CORMed); Hangzhou, China; Department of Engineering Mechanics, Zhejiang University; Hangzhou, 310027, China; School of Materials Science and Engineering, Nanyang Technological University; Singapore, 639798, Singapore; Beijing Life Science Academy; Beijing, 102200, China; Center of Regenerative and Aging Medicine, the Fourth Affiliated Hospital of School of Medicine, and International School of Medicine, International Institutes of Medicine, Zhejiang University; Yiwu, 322000, China; School of Materials Science and Engineering, Zhejiang University; Hangzhou, 310058, China; Department of Orthopedics, Shanxi Key Laboratory of Bone and Soft Tissue Injury Repair, Second Hospital of Shanxi Medical University; Taiyuan, China; Mechano-X Institute, Applied Mechanics Laboratory, Department of Engineering Mechanics, Tsinghua University; Beijing, 100084, China

**Author notes:** These authors contributed equally to this work (Wenyue Li, Xiaozhao Wang, Renwei Mao, Dong Li).

## Abstract

Joining heterogeneous materials in engineered structures remains a daunting challenge because of stress concentration, often resulting in unexpected failures^1,2^. Studying the structures in organisms that evolved for centuries provides valuable insights that can be instrumental in addressing this mechanical challenge^3–5^. The human meniscus root-bone interface is a remarkable example known for its exceptional fatigue resistance, toughness and interfacial adhesion properties throughout its lifespan^6–8^. We studied the multiscale graded mineralization structure designs within the 30-micron soft-hard interface at the root-bone junction and examined its toughening mechanisms. This graded interface with interdigitated structures and exponential modulus increase exhibits a phase transition from amorphous calcium phosphate (ACP) to gradually matured hydroxyapatite (HAP) crystals, mediated by location-specific distributed biomolecules. In coordination with collagen fibril deformation and reorientation, ACP particles debond with collagen and slide to new positions which enable frictional energy dissipation, and HAP particles arrest cracks. The mineral in transforming phases work synergistically to provide interfacial toughening. The presented biointerface model exemplifies human musculoskeletal system’s adaptations to mechanical requirements, offering a blueprint for developing tough interfaces in broad applications.

## Introduction

Population aging presents significant challenges to the global healthcare system, particularly regarding disease, disability, and cognitive decline. As we enter an era of increased longevity, it becomes crucial to maintain the functionality of tissues and organs over time, or to provide convincing solutions to repairing and regenerating the defected ones. In line with such demand, constructing durable interfaces between materials with differing mechanical properties is vital for extending the lifespan of biomedical devices such as implants^9^, biosensors^10^ and tissue engineering constructs^11^. However, joining dissimilar materials is vulnerable to stress concentration that commonly induces premature interface failure^1^. Nature offers valuable inspiration for interfacial bonding solutions, with intricate structures which have evolved to function in various harsh loading scenarios, such as the enthesis, squid beak, and diabolical ironclad beetle^12^ showcasing multiscale gradients in structure, composition, and mechanics that mitigate localized stress. While we have learned from the extrinsic mechanisms of these natural structures, the toughening and anti-fatigue mechanisms in humans’ native tissues remain to be uncovered in endeavors to maintaining or regenerating human tissues, and to deepen our understanding of interface design principles.

The human meniscus root to bone presents as an excellent model of interfacial toughening^8^. This junction tissue anchors the meniscus body to the tibial bone, protects the knee joint by redistributing loads and converting axial loads into hoop stresses^6,7^ (Fig. 1A). Meniscus root bears tension up to 200 N with a 20 mm^2^ cross-sectional area^13,14^, in which the stress is comparable to using a single hand to lift a five-ton truck. The demanding mechanical loading regime drives the root-bone interface to evolve into an extremely tough, fatigue-resistant soft-hard transitional structure. It consists of a ligamentous root (LR), fibrocartilage (FC), mineralized fibrocartilage (MFC) and bone (B) in histological classification (Fig. 1B,C)^15^, with tissue modulus varied gently from 2MPa in LR to 10 MPa in FC, while exponentially increased to over 1 GPa within the 30-μm soft-hard (S-H) interface (Fig. 1D, Fig. S2). Despite the sharp modulus transition at the S-H interface, the root-bone interface can achieve excellent anti-fatigue adhesion and toughness^14,16,17^, exhibiting high durability in millions of loading cycles annually. Although previous studies have attributed the interface’s resilience to its graded organization and extracellular matrix (ECM) compositions consisting of organic and inorganic mineral aggregates^15^, the specific contributions of these gradients to durability and the associated toughening mechanisms remain less understood. Complex assembly and phase transition of mineral aggregates play a dominant role in determining tissue mechanical properties in numerous natural materials, including interfaces^5,18,19^. The mantis shrimp’s dactyl club obtains exceptional strength from amorphization and dislocation of hydroxyapatite granules^20^; amorphous intergranular phase between hydroxyapatite improves the enamel’s hardness and resistance to acid^21,22^; the nanoscale gradient in mineralization at the osteochondral interface effectively prevents deformation localization^5^. These findings highlight the importance of mineral phase and its distinct assemblies with surrounding matrix to the interfacial robustness and inspire us to systematically study the detailed matrix-mineral assemblies and their unique fatigue-resistant and toughening mechanisms at the root-bone interface.

**Fig. 1.**
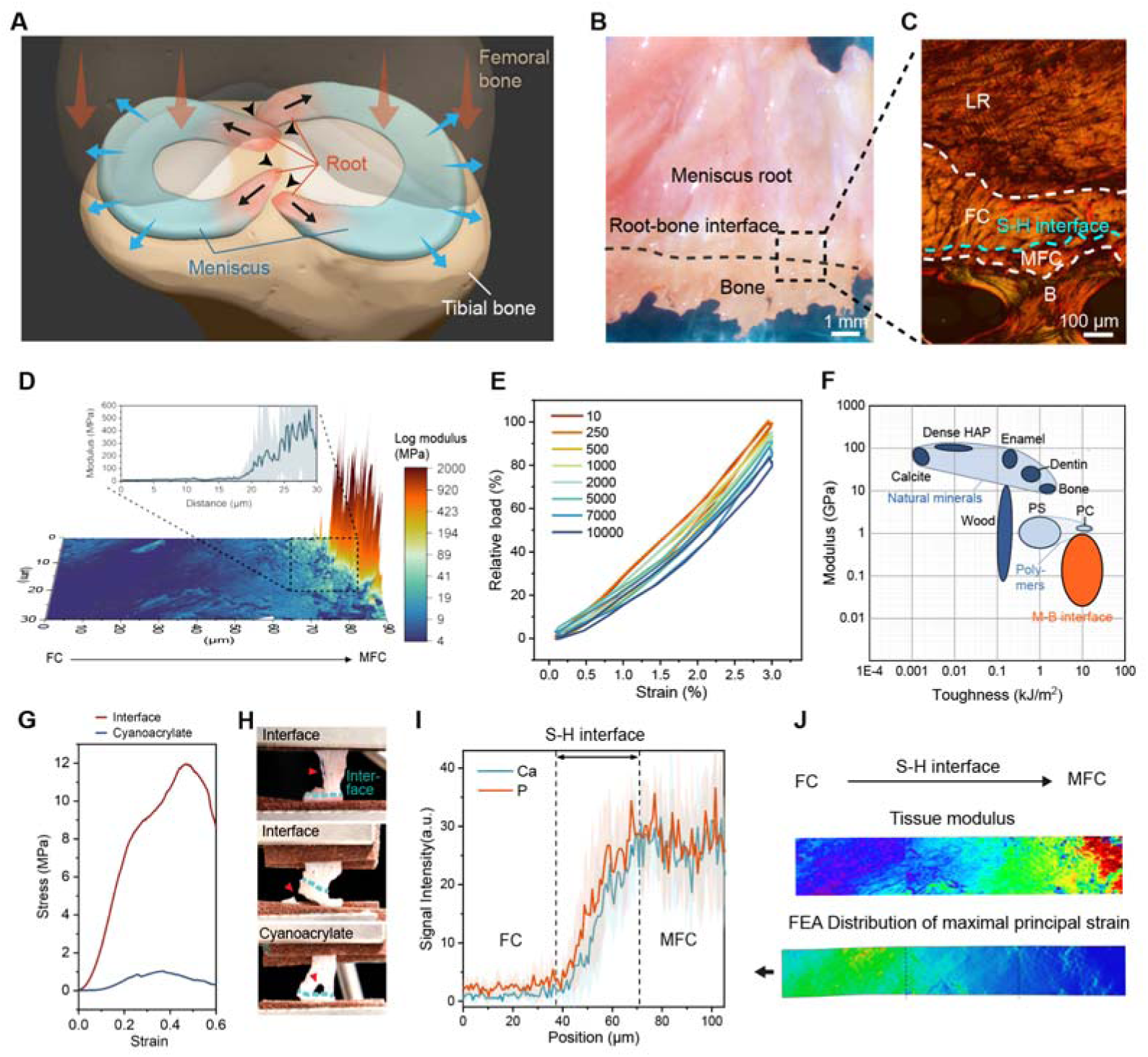
Overview of the meniscus root-bone interface tissue structure and mechanical properties. (**A**) Schematic illustration of the loading environment of meniscus roots. (**B**) Optical image of the root-bone interface. (**C**) Picrosirius red staining of the root-bone interface revealing ligamentous root (LR), fibrocartilage (FC, 200-300 μm), mineralized fibrocartilage (MFC, 50-100 μm), and bone (B). (**D**) Young’s modulus data of tissue obtained using AFM. The enlarged view showed the modulus change trendline at the S-H interface. (**E**) Relative load of multiple cycles during the fatigue test of root-bone tissue. (**F**) Ashby plot of modulus and toughness of natural and synthetic materials. (**G**) Interface adhesion tests showing the interface adhesion strength of native tissue and cyanoacrylate. (**H**) Optical images of unloaded and failed samples in interface adhesion test. Failure appeared within the root or bone region for the native interfacial tissue, yet at the junction part for the cyanoacrylate-adhered interface. (**I**) EDS line scan showing calcium (Ca) and phosphorous (P) intensity across the S-H interface. (**J**) FEA results of strain distribution in a tensioned S-H interface sample, with the modulus data from AFM measurement. Abbreviations: S-H interface, soft-hard interface; PS, polystyrene; PC, polycarbonate; EDS, Energy Dispersive X-ray scanning; FEA, finite element analysis.

### The meniscus root-bone interface demonstrates exceptional mechanical performance

To quantify the anti-fatigue and load-bearing properties of the root-bone interface, we conducted targeted mechanical tests on porcine meniscus root-bone tissue, spanning from LR to bone. The tissue displayed no apparent sign of fatigue after 10,000 cycles of physiological loading, verifying its superior fatigue resistance (Fig. 1E; Fig. S3). Fracture toughness measured using interfacial tissue reaches 12 kJ/m^2^, surpassing that of most biological materials and synthetic polymers as represented in the Ashby plot (Fig. 1F)^23,24^. Furthermore, the root-bone interface outperformed the most commonly used commercial adhesive cyanoacrylate in adhesion strength (Fig. 1G); Unlike cyanoacrylate-treated interfaces which failed at adhesion site, the native interfacial tissue failed at root or bone tissue (Fig. 1H), indicating outstanding adhesive properties. The outstanding mechanical performance of the root-bone interface implicates that the potential stress concentration problem was effectively resolved at the soft-hard (S-H) interface spanning ~30 μm, as observed via Energy Dispersive X-ray scanning (EDS) (Fig. 1I). The intrinsic modulus transition pattern turns out to be desirable for stress concentration mitigation, as monitored in Finite Element Analysis (FEA) (Fig. 1J; Fig. S4; Movie S1), indicating that the native modulus gradient enables proper interfacial load redistribution, which is essential for preventing fatigue^25^.

### Minerals exhibit an elaborate gradient across the meniscus root-bone interface

As the determining factor of tissue stiffness, mineralization at the S-H interface merits thorough examination^5,18,19^. Focused ion beam-scanning electron microscope (FIB-SEM) analysis reveals three distinct mineral morphologies at the soft-hard (S-H) interface as mineral content increases (Fig. 2A-B): spherical particles (region i; Fig. 2C; Movie S2), fused morphology (region ii, Fig. 2D), and dense space-filling state (region iii; Fig. 2E). SEM, surface topography mapping (Fig. 2F-K), as well as an AFM modulus map which revealed a granular increase of modulus (Fig. S5), all reflected similar mineral morphological transformations^26^. The spatial relationship between minerals and collagen fibrils varies across the interface, with extrafibrillar distribution in region i transitioning to intrafibrillar occupation of minerals in region ii and iii (Fig. 2C-E; Fig. S5 C-E; Fig. S6), which resulted in an incremental loss of fibril structural integrity (Fig. S7). Interestingly, closer inspection of region i shows that extrafibrillar minerals are in close contact with fibrils (Fig. S7G-J), implicating potential interactions at a smaller scale. The three-step morphological transition can also be visualized on transmission electron microscopy (TEM) images (Fig. S8). The subtle changes in distribution and morphology in the gradient mineralization zone (Table S2) suggest underlying compositional transitions of mineral aggregates^27^.

**Fig. 2.**
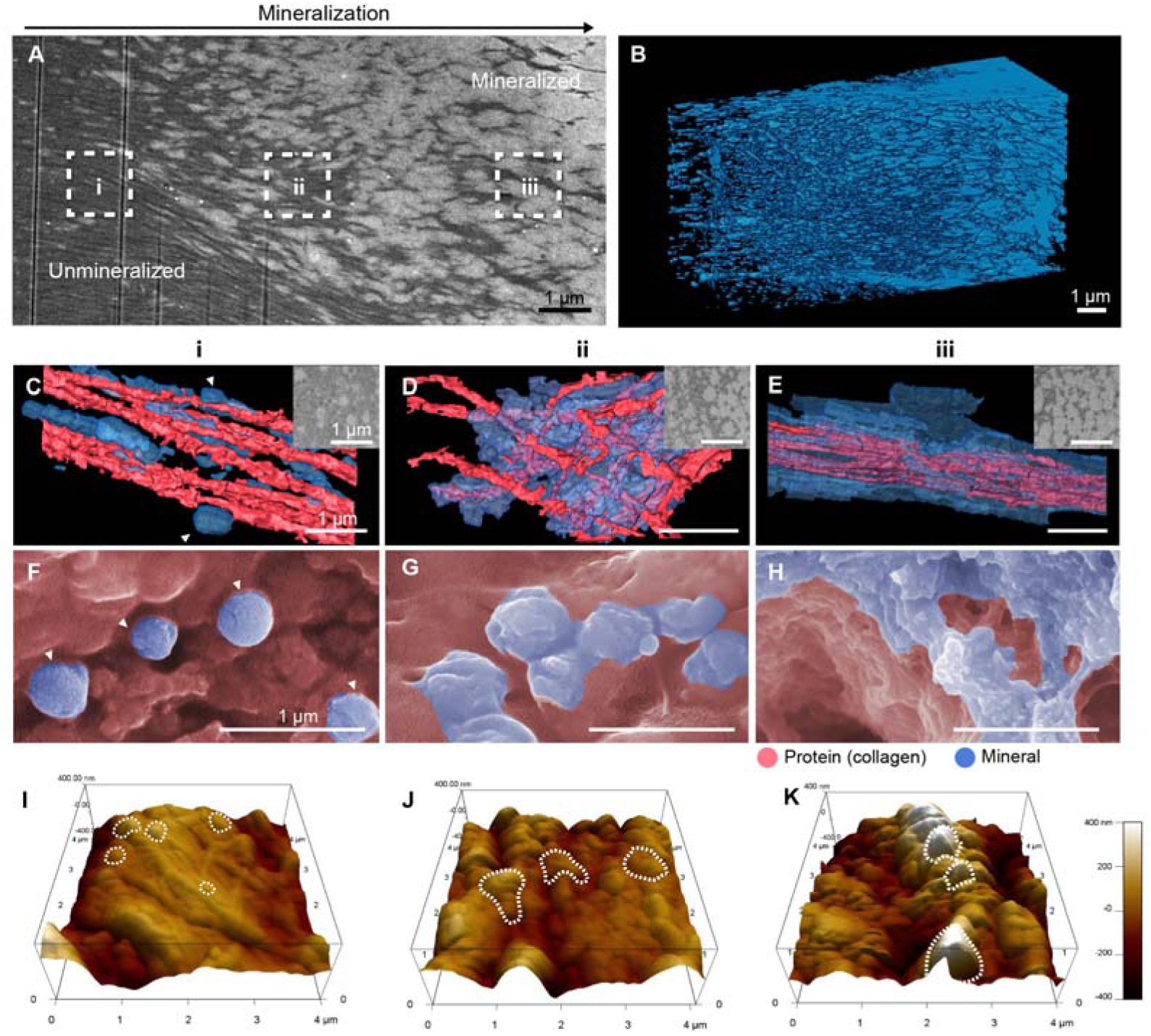
Minerals exhibit an elaborate pattern transition across the S-H interface of meniscus root. (**A**) Longitudinal view of the soft-hard (S-H) interface. FIB-SEM data at the root insertion which illustrates the increasing density of mineralization of fibrocartilage. Outlined areas (i,ii,iii) in white dash line are representatives of the transitional patterns of minerals. (**B**) FIB-SEM 3D reconstruction of minerals. (**C-E**) 3D reconstruction of 3 transitional regions in (a): (i) spherical minerals in the extrafibrillar space; (ii) irregular minerals; and (iii) dense minerals in the extrafibrillar and intrafibrillar space. Insets show 2D cross-sectional views of corresponding regions. (**F-H**) Colored SEM images of region i-iii showing similar patterns in mineral morphology. Image originates from SEM with backscattered signals. (**I-K**) 3D topographical maps of region i-iii presenting the transition from spherical particles to typical mature mineral platelets on sample surface.

### Transformation of the mineral phase across the meniscus root-bone interface

To elucidate the compositional nature of the mineralization gradient, the S-H interface was further observed via Raman spectroscopy, spherical aberration-corrected transmission electron TEM and Cryo-TEM. Raman spectra and mapping data show that, from the unmineralized to the mineralized region, the S-H interface exhibits an increasing signal of hydroxyapatite (HAP) band relative to proteins and increasing HAP crystallinity, along with el6vated intensity of carbonate and a lower rate of carbonate substitution (Fig. S9; band assignments listed in Table S3), indicating gradual maturation of HAP at the S-H interface^28^. We also observed a shift of the HAP signal to the lower spectrum at the frontier of the S-H interface, probably indicating the presence of an immature mineral or amorphous calcium phosphate (ACP) phase^29^.Similar signal change was also observed in porcine S-H interface (Fig. S10). This could be validated by Stimulated Raman Scattering (SRS) with higher spatial and spectra resolution^30^, which shows a clear conversion of ACP to HAP at the S-H interface (Fig. 3A; Fig. S11). TEM images also revealed transition from amorphous state to gradually crystalline state (Fig. 3B). Given the unstable nature of ACP, we further examined the mineral phase and structures using cryo-TEM, and the results demonstrate a non-crystalline characteristic of sphere and chain-like aggregates (stage i) present at the frontier of the S-H interface (Fig. 3C-D), corresponding to the spherical morphology in region i in Figure 2. At stage ii, mineral crystals emerged and changed from rod-like to platelet-like shape with increasing density. Deeper into mineralized region, mineral deposits transformed to a mature lacy pattern (stage iii)^18^, collectively presenting a three-stage subtle transition of mineral assembly patterns at the S-H interface.

**Fig. 3.**
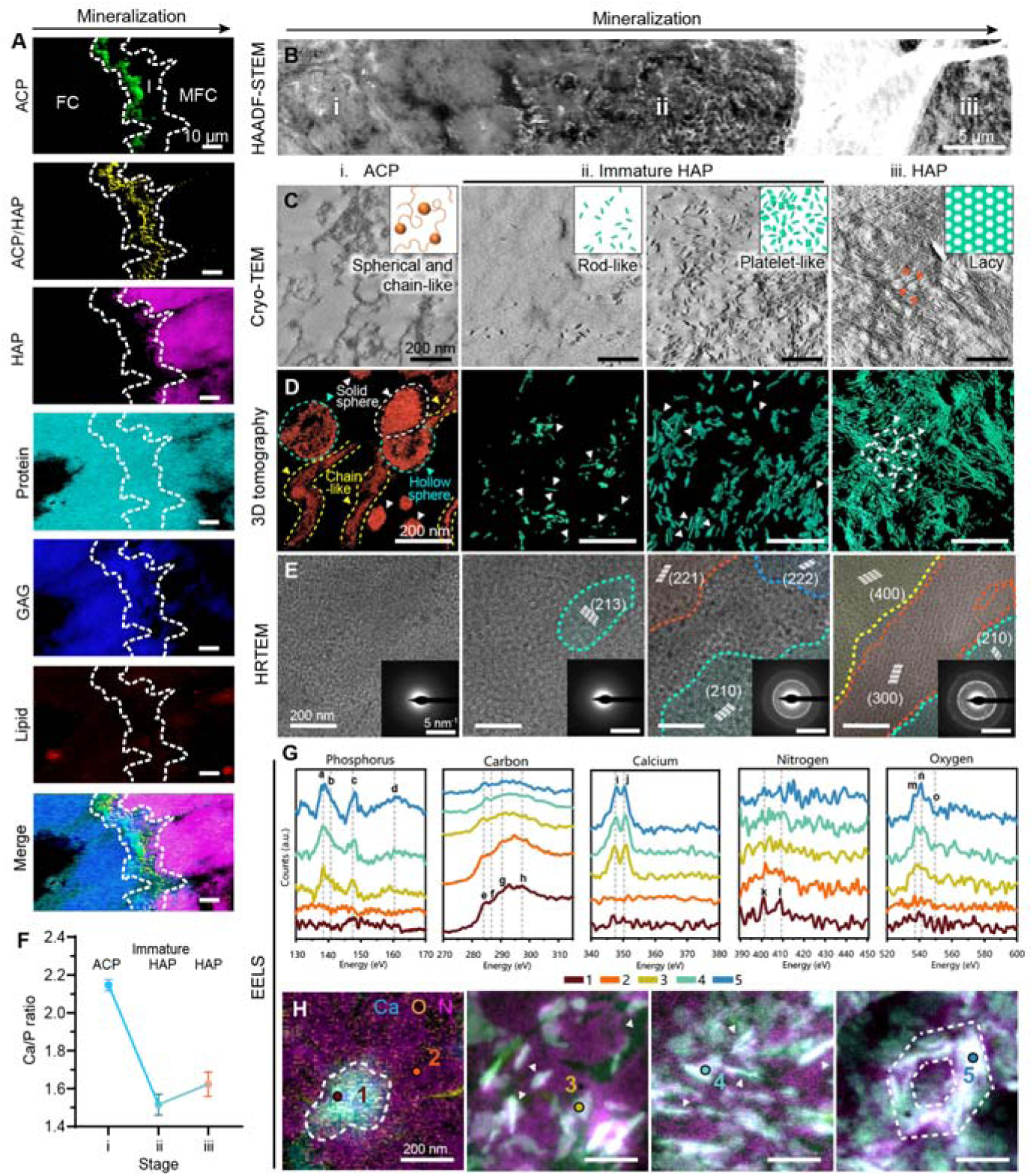
Phase transformation of minerals across the meniscus root-bone interface. (**A**) SRS intensity imaging of the S-H interface. (**B**) An HAADF-STEM snapshot of the entire S-H interface showing the mineral phase transition. (**C-D**) Cryo-TEM images and 3D tomography of minerals across the S-H interface (configurations summarized in inlets). Minerals at stage i show hollow sphere, solid sphere and chain structures. Stage ii minerals transitioned from rod-like to platelet-like morphology with increasing HAP density. Stage iii minerals were in thinner platelet shape and aggregating into a lacy pattern. (**E**) HRTEM and SAED results of different stages. HRTEM demonstrates gradual increase of HAP lattice. SAED patterns in inlets reveal the transformation from broad diffuse ring to clear diffraction spots and thin rings. (**F**) Ca/P molar ratio of minerals analyzed through EDS. (**G-H**) EEL spectra and intensity mapping. Representative spectra at positions denoted in (H) are shown in (G). Phosphorus L_2,3_ edge, Carbon K edge Calcium L_2,3_ edge, Nitrogen K edge and Oxygen K edge heatmaps were generated through a power law model after normalization. Abbreviations: SRS, stimulated Raman scattering; ACP, amorphous calcium phosphate; HAP, hydroxyapatite; GAG, glycosaminoglycan; HAADF-STEM, high-angle annular dark field scanning TEM; HRTEM, high resolution TEM; SAED, selected area diffraction; EEL, electron energy loss.

Additionally, the mineral crystals exhibited size, flatness and aspect ratio variation from stage ii to iii; in particular, crystals in stage iii displayed morphological features of bone minerals (Fig. S12)^18^. High-resolution TEM (HRTEM) and diffraction patterns (Fig. 3E; Fig. S13) showed that stage i has poor crystallinity, which increased in stage ii and iii, indicating gradual maturation of minerals. Examination of the Ca/P ratio across stages (2.146±0.2946 (i), 1.516 ±0.05499 (ii) and 1.623±06386 (iii)) further supported the presence of ACP in stage i and progressive maturation of HAP from ii to iii^31^ (Fig. 3F, Fig. S14). The elaborate mineral assembly transformation in phase and crystallinity at the S-H interface presents a complete evolution of minerals. This gradient in mineral phase could serve as a buffer zone and enhance adhesion between tissues with disparate mechanical properties, analogous to the amorphous layer between HAP platelets and organic matrix in nacre^32^. Regardless of the unstable nature of immature mineral and its tendency to convert into HAP^29,33,34^, ACP was identified in adult human interfacial tissue, implicating potential mediating mechanisms of local mineral phase transition.

Next, electron energy loss spectroscopy (EELS) was utilized to examine the distinct chemical environment of varied minerals at the S-H interface. Augmentation of Ca and P signals from stage i to iii indicated increased inorganic mineral content, suggesting a three-staged transition tendency of minerals; meanwhile, N and C peaks declined gradually, which indicates a decrease in proportion of organic components (Fig. 3G-H, Fig. S15). In particular, the peak of carbonyl group (peak f) and nitrogen K edge (peak k, l) were enriched only in front stages (i, ii) as shown in Figure 3G, suggesting a positional correlation between immature minerals and organic matrix (proteins)^31^, which may be the potential mechanism for immature minerals maintaining stability.

### Biomacromolecules mediate phase transformation of minerals at the meniscus root-bone interface

The existence of ACP *in vivo* relies on stabilizing molecules, especially proteins^35^ colocalized with the ACP phase in the S-H interface in EELS data (Fig. 3F). To identify potential protein stabilizers, liquid chromatography−tandem mass spectrometry (LC-MS/MS) was conducted on meniscus root, root-bone interface and bone sections (Fig. 4A). Manual dissection might introduce remnants of root or bone tissue at the interface, potentially obscuring protein expression differences. Among the 1,026 proteins detected at the root-bone interface, 40 exhibited significantly higher expression levels compared to bone, and 16 showed higher expression compared to root (>twofold difference) (Fig. S16, Supplementary data 1). Enriched proteins were primarily associated with ECM constituents and cartilage homeostasis (Fig. 4B, Fig. S17), such as collagen II, aggrecan and biglycan, likely reflecting the presence of fibrocartilaginous tissue at the interface and its role in withstanding shear and compressive stress^4,11^. Notably, matrix Gla protein (MGP), a mineralization inhibitor^36^, was highly expressed at the interface compared to root tissue (Fig. 4C). MGP exhibited a location-specific distribution at the mineralization front at the S-H interface, as shown by immunofluorescent staining (Fig. 4D), and correlated with calcium which was indicative of mineralization degree (Fig. 4E).

**Fig. 4.**
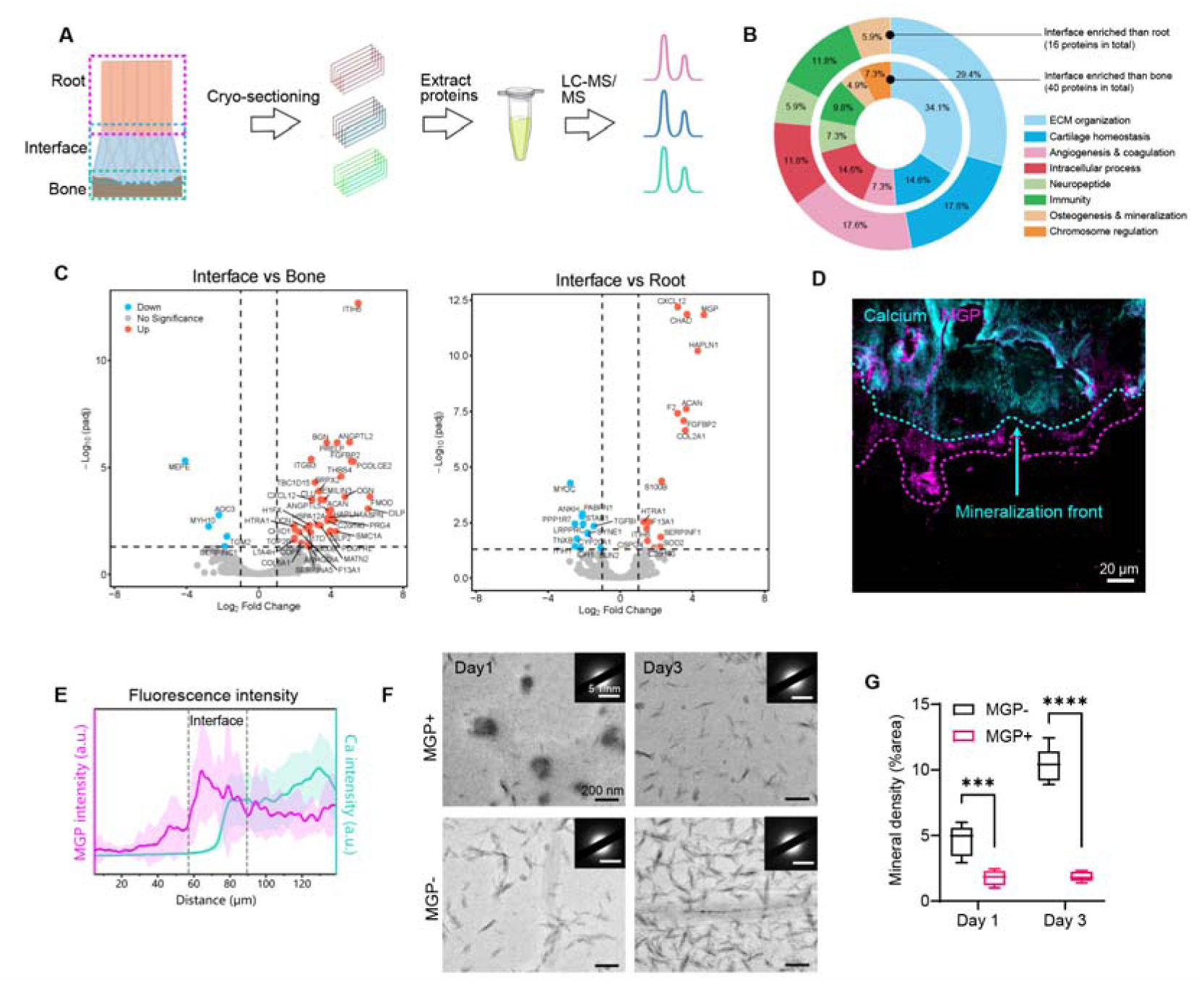
Root-bone interface-abundant proteins are important contributors to interface mechanics and mineralization. (**A**) Schematic illustration of the experimental procedure of proteomic analysis. (**B**) Classification of interface-enriched proteins based on functions described in Genecards Database. (**C**) Volcano plots of protein expression differences between interface and bone, and interface and root. Proteins with statistically significant differences in expression were colored in orange (significantly higher in interface) and blue (significantly lower in interface) (*p* adjusted value < 0.05, fold change > 2). (**D**) Immunofluorescent staining of MGP at the S-H interface. (**E**) MGP protein expression profile across the S-H interface quantified using fluorescent signal intensity which increased in the mineralized tissue region and S-H interface (denoted by dashed line). (**F**) Functional verification of MGP’s role as a mineralization inhibitor through *in vitro* mineralization using nano-ACP. TEM images showed the increase in HAP crystals in the non-treated group was less than in the MGP-treated group on day 1 and day 3. Inlets revealed diffraction patterns of associated areas. (**G**) Quantification of HAP crystal density in TEM snapshots (n=5) of MGP-treated and non-treated groups. Statistical significance is illustrated as: *p ≤ 0.05; **p ≤ 0.01; ***p ≤ 0.001. Abbreviations: LC-MS/MS, liquid chromatography−tandem mass spectrometry; MGP, matrix Gla protein.

Previous reports suggest that MGP has a high affinity for HAP crystals and hinder mineralization of HAP^37^. We hypothesize that MGP could serve as a crucial mediator in the mineral phase transition from ACP and immature HAP to mature HAP at the S-H interface. To address this hypothesis, an *in vitro* mineralization experiment was conducted using nano-ACP (details described in methods), which would transform into HAP crystals at 37 °C in 3 days. Remarkably, MGP-treated nano-ACP resulted in the formation of spherical amorphous aggregates of Ca and P at day 1(Fig. S18), followed by delayed crystallization and reduced HAP density at day 3 (Fig. 4F-G), suggesting that MGP acts as an important ACP stabilizer at the interface, maintaining the dynamic balance between unmineralized and mineralized tissue. In addition, the AFM adhesion map illustrated a local increase in adhesion force at the S-H interface area (Fig. S19), correlating with small leucine-rich proteoglycans such as biglycan, fibromodulin and decorin, which were highly enriched in interfacial tissue (Supplementary data 1)^38^. These proteoglycans are important regulators of tissue mineralization and collagen fibril sliding^39,40^, which could enhance toughness^41^ and facilitate at stable transition between dissimilar tissues^4^. Collectively, the biomolecules enriched at the S-H interface could regulate mineral phase transformation and shape tissue mechanics.

### Synergy of mineral and collagenous matrix toughens the interface

To elucidate the significance of hierarchical structural and compositional transition in interface toughening, we investigated the mechanical response of interfacial tissue from the tissue level to the molecular level. Tensile loading was applied to fluorescently labelled thin root-bone specimens, and local displacement was visualized under a microscope (Fig. S20). Generally, the root-bone tissue exhibited fiber straightening and reorganization in soft tissue, followed by involvement of the interface and subsequent soft tissue failure (Fig. 5A). The LR and FC region experienced deformation at low level of strain, while the interface region deformed only when strain exceeded 15% (Fig. 5B, Fig. S21). This agreed with FEA results, in suggesting that unmineralized tissue dominates mechanical response due to greater deformability (Fig. 1J), which could lower the risk of brittle failures in the S-H interface and mineralized tissue caused by excessive straining, laying the basis for superior fatigue resistance. To study potential strategies of the S-H interface in dealing with high stress and flaws, a crack was generated at the interface area. Upon stress, crack deflection toward soft tissue or occasionally mineralized region was observed (Fig. S22, Fig. S23A). This mechanism emerges as another inherent response to prevent defects in interfacial tissue; meanwhile, the cracks would be terminated in soft tissue through blunting^42^ (Fig. S22B) or arrested by stiff mineral aggregates deposited in the matrix. Delving closer in the notch zone (Fig. S23B, E), crack blunting was achieved by collagen fibril bridging, rotating and re-aligning, accompanied with the breakage of interfibrillar bridges (Fig. S23C, F), exhibiting distinct morphology compared to a cut-made notch (Fig. S23D). The S-H interface was rarely affected by the notch zone, but a stress-concentrated plastic zone forming ahead of the notch could involve it and lead to microscale flaws (Fig. S23G). Various energy dissipation mechanisms were employed at the S-H interface to minimize stress: unmineralized collagen fibrils straightened and slid to dissipate energy (Fig. S23H); in the graded mineralized region, partially interfibrillar mineralization endowed efficient force transfer from soft collagen fibrils to hard mineral aggregates (Fig. S23I, J; Fig. S24); the fully mineralized region’s high stiffness resisted further crack development (Fig. S23G). Furthermore, consistent with post-straining SEM results (Fig. 5C), *in-situ* SEM observation of root-bone specimens under tensile stress showed movement of interfibrillar mineral aggregates relative to matrix at the S-H interface during straining (Fig. 5D, Fig. S25), akin to the bone toughening mechanism—friction generated between mineral particles and the matrix could dissipate energy as heat^43^. On a finer scale, electrostatic attraction between the immature mineral aggregates and collagen^44^ revealed by Molecular Dynamics (MD) modelling also contributed to energy dissipation and is instrumental for interface adhesion (Fig. 5E)^45^. Destruction and re-establishment of electrostatic interaction could occur simultaneously, constantly toughening the interface at nanoscale Additionally, breakage of interfibrillar bridges (Fig. 5F) further contributed to collagenous matrix toughening^46^.

**Fig. 5.**
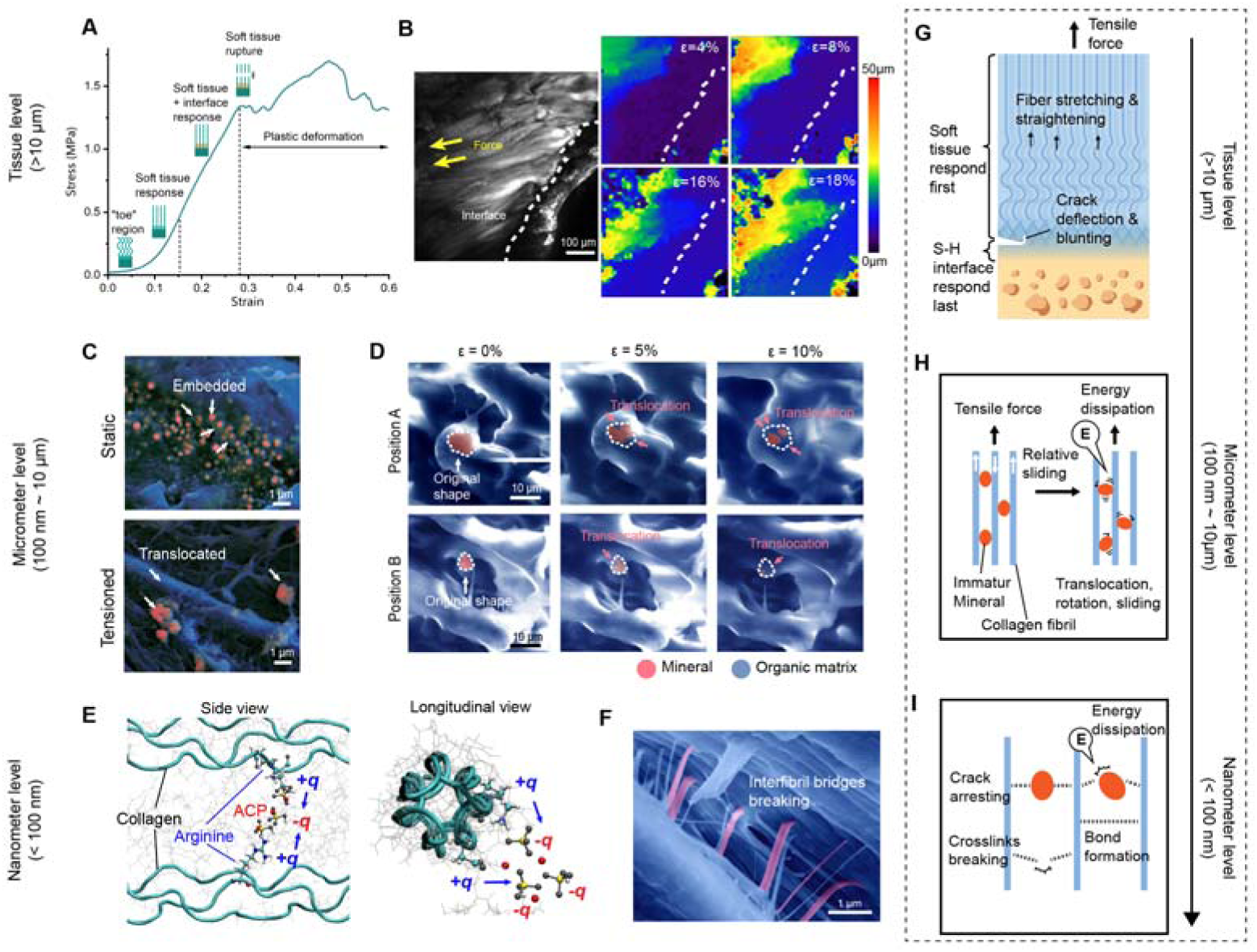
Multiscale toughening mechanisms of the meniscus root-bone interface. (**A**) Typical stress-strain curve of the root-bone interface under tensile stress and four stages of tissue response. Crimped fibers in the soft tissue straightens in the “toe” region; as the curve becomes linear, soft tissue continues to respond to stress in a linear fashion, while the S-H interface responds lately; failure of tissue generally occurs at soft tissue during plastic deformation. (**B**) *In-situ* observation of root-bone tissue response to tensile stress. Maps of displacement magnitude of tissue under different strain (ε) levels are shown. **(C**) SEM images demonstrate the interfibrillar spherical minerals underwent translocation from their original positions at the static state upon stress. (**D**) Observation of spherical minerals through *in-situ* SEM. (**E**) Snapshots of molecular dynamic modelling of ACP-collagen interaction. Arginine and ACP interact because of electrostatic attraction. (**F**) SEM image showing fibril tearing as a result of interfibrillar bridges breaking. (**G**) Schematic illustration of tissue level toughening. (**H**) Schematic description of toughening mechanisms at the microscale: mineral sliding along fibrils together contributes to energy dissipation. (**I**) Graphic illustration of nanometer-level energy dissipation through the breakage of molecular bonds.

The native phase transition of minerals is significant for the interfacial mechanical features, as illustrated by over-mineralization and demineralization which resulted in reduced toughness and strength (Fig. S26A-C). The bone tissue of demineralized samples resembled elastic materials that exhibited enhanced ductility but with much lower strength, while over-mineralized samples had higher rate of brittle fracture at the bone; interestingly, the failure mode of both treatments was avulsion from bone followed by interfacial delamination in perpendicular direction, indicating interfacial weakening compared to the untreated samples (Fig. S26D).

Collectively, we have demonstrated multiple toughening mechanisms achieved by collagen-based matrix and phase-transforming minerals at three levels. ① At the tissue level (>10 μm), deformation is dominated by soft tissue, including LR and FC, barely involving the S-H interface. Cracks occurring at the S-H interface tend to be redirected to non-interfacial tissues to prevent further flaws at the interface area (Fig. 5G). ② At the micro-nano level (100 nm ~ 10 μm) within the S-H interface region, energy can be dissipated through collagen fibril straightening, re-aligning, rotating as well as stress-induced frictional sliding and crack arresting of the mineral aggregates (Fig. 5H). ③ At the nanometer level (<10 nm), breakage of fibril-mineral and interfibrillar bonds facilitates energy dissipation and improves toughness (Fig. 5I).

### Phase transformation toughens biointerface composites

To further test the potential benefits of mineral phase transition in meniscus root-bone interfaces, we constructed artificial root-bone composites composed of freeze-casted silk fibroin methacrylate (silMA) with aligned fibers mineralized with a suspension of ACP and HAP nanoparticles (Fig. S27A). Mineralization reinforced the fibrous structure by incorporating into the matrix (Fig. S27B) and generated soft-hard interfaces with or without an ACP interlayer. Results show that the addition of ACP interlayer significantly increased the tensile modulus (5.92±0.57 MPa) compared to silMA only (3.04±0.73MPa) and HAP+ root (3.28±0.4) as well as toughness (NC: 0.121±0.025 MJ/m^3^; ACP: 0.15±0.030 MJ/m^3^; HAP: 0.118±0.037 MJ/m^3^; ACP+HAP: 0.285±0.073 MJ/m^3^) (Fig. S27C, D). The failure modes of mineralized structures also indicate the importance of ACP-HAP transition at an interface with dissimilar material properties (Fig. S28). The presence of the ACP interlayer may help reduce the interfacial energy between soft and hard regions by replacing a single energy-costly interface with two interfaces with lower energy to improve the mechanical performance of the heterogeneous root^32,47^. Therefore, we demonstrated the significance of ACP-HAP transformation for mechanical properties of an artificial engineered root-bone interface, providing inspirations for reinforcement of soft-hard materials.

## Discussion

Our study has uncovered interfacial features in human meniscus root-bone interfaces that ensure a stable connection between mechanics-dissimilar tissues and efficient force transfer to maintain knee stability. We provide experimental observations and simulation results confirming the multiscale toughening mechanism at the S-H interface, especially within the ultra-thin, 30 μm gradient mineralization region (Fig. 6). Macroscopically, the root-bone interface exhibits interdigitated structures and graded stiffness that help distribute load uniformly to protect the tissue. Further structural and mechanical characterization highlights the benefits of the elaborate mineral assemblies, including phase transition from ACP to HAP and three-stage mineralization in the collagenous matrix over the S-H interface. The high deformability of the unmineralized soft matrix region provides crack deflection and blunting capability to the superior interfacial toughness^42^, while the fully-mineralized region, where mineral content reaches 60-70%^48^, undergoes little deformation and safeguards against crack propagation. Notably, the gradient mineralization area possesses immature mineral particles in spherical shape located between collagen fibrils. These immature minerals could undergo sliding in response to tensile stress on the interface, resulting in energy dissipation in the manner of frictional energy and molecular bond-breaking energy at the mineral-collagen interface. These multiscale organic-inorganic assembling structures work synergistically to provide toughening mechanisms, thus avoiding failure of the S-H interface (summarized in Movie S3).

**Fig. 6.**
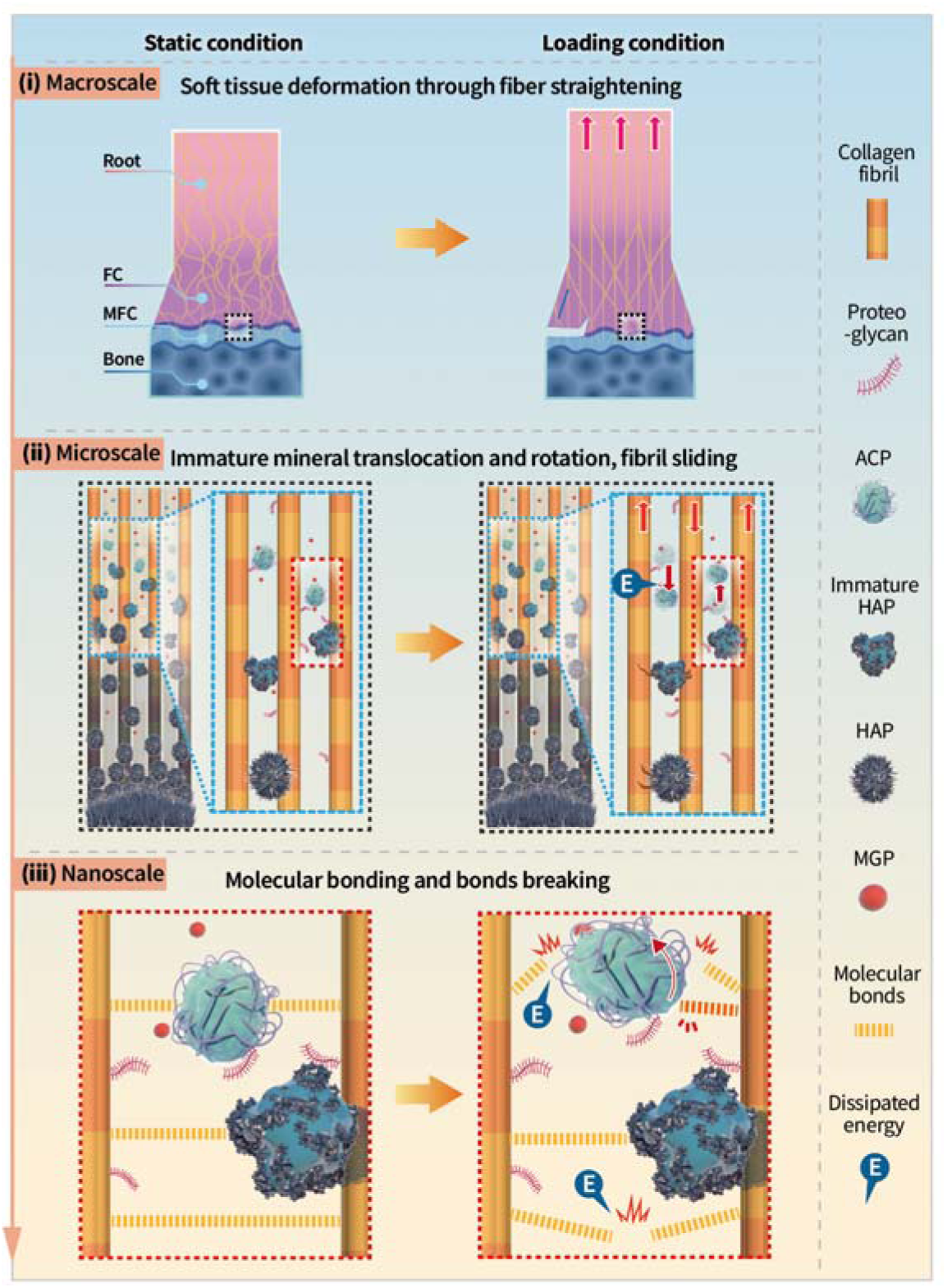
Schematic summary depicting toughening mechanisms of the meniscus root-bone interface. The left panel illustrates the hierarchical structure of the meniscus root-bone interface under static condition. From macroscale to nanoscale: histological zones of the root-bone interface; transformation of mineral aggregates from ACP, immature HAP to HAP alongside mineralization of collagen fibrils; ACP, immature HAP and collagen form molecular bonds with collagen fibrils, while MGP protein and proteoglycans interact with the mineral aggregates and collagen to mediate mineralization. The right panel demonstrates the multiscale mechanical response upon mechanical loading. From macroscale to nanoscale: soft tissue responds to stress first via fiber stretching and straightening while the interface responds last, and cracks would be deflected or blunted; the ACP and immature aggregates dissipate stress (denoted by “E”) through sliding in the interfibrillar space, and crack would be arrested by stiff mineral aggregates; bonds among mineral aggregates and collagen could break for energy dissipation, and new bonds formation could occur simultaneously. Abbreviations: LR, ligamentous root; FC, fibrocartilage; MFC; mineralized fibrocartilage; ACP, amorphous calcium phosphate; HAP, hydroxyapatite; MGP, matrix gla protein.

Parametric studies of the biocomposite adhesive and artificial root-bone structures demonstrates an immediate benefit of mineral phase transition at the interface, providing increased toughness and substantial strength of scaffolds^22,48,49^. Viewed as a polymer-based material reinforced by nanoparticles, the interfacial tissue exhibits high plasticity, influenced by factors like the composition of nanoparticles, polymer characteristics, their distribution and density, and their interplay^50^. Looking forward, by combining advanced synthesis and manufacturing technologies with the design principles from human complex structures, a bottom-up strategy to build multiscale composite interface materials with broad applications, including brain-machine interfaces, wearable devices, implanted biosensors as well as anti-fatigue soft-hard structures, could be achieved.

## Methods

### Human sample preparation

Workflow of processing samples and subsequent analysis are presented on Fig.S1. Knee samples were collected with written informed consent from patients undergoing amputation due to trauma (n=5). None of the donors were engaged in athletic professions, and all exhibited normal knee joint conditions. All specimens were obtained and used according to guidelines approved by the Second Hospital of Shanxi Medical University Ethics Committee (Ethical NO. 2019YX260). Meniscus root tissues including part of the meniscus, the ligamentous root, and subchondral bone were dissected from the tibial plateau and washed carefully with cold phosphate-buffered saline (PBS). Samples were split into 3-5 pieces (each piece have complete interfacial transition from ligamentous root to bone), among which one piece was fixed and decalcified for histological analysis, and the other pieces were cryo-sectioned into 5 mm and 15, 30, 50, 100, 200 μm-thick specimens in the longitudinal direction and stored at −80 °C until being used for experiments.

Donor and sample information is presented in Table S1. Meniscus roots in four positions can be found in one knee: lateral anterior, lateral posterior, medial anterior, medial posterior roots. Clinical data demonstrated that the medial posterior root tear is the most prevalent root tear (accounting for 52.1% of root tears)^51^. Injury at medial posterior root leads to aberrant knee biomechanics which can be equivalent to total meniscectomy^52^. Due to limited number of human samples and the requirement to eliminate potential variance of anterior or posterior, medial, or lateral meniscus root samples, medial posterior root—the site with the most clinical concern, was chosen for analysis and results presenting while other roots were used mainly for optimization of sample preparation and experimental procedures.

### Histological staining

Meniscus root samples for histological analysis were fixed for 24-48 hours and decalcified for two months, dehydrated and embedded in paraffin. After obtaining 10-μm paraffin sections using Leica RM2235 Paraffin Microtome (Leica, Hamburg, Germany), Hematoxylin and Eosin staining, Picrosirius Red staining were conducted on the sections following standard protocols and observed through polarized microscopy.

### Fatigue test

For fatigue test of meniscus root-bone interfacial tissue, a porcine stifle joint was collected immediately after sacrifice and kept on ice before dissection and isolation of meniscus root-bone interfacial tissue specimen. Porcine meniscus root samples were used because of the requirement for bioactivity in mechanical characterization and limited quantity of human samples; porcine meniscus is frequently utilized to model human meniscus due to their similarities.^53,54^ To ensure cell viability during the test, the loading device (MTS Criterion 40) was framed in a custom-made constant temperature and humidity chamber in which 37 °C and 90 % humidity (humidifier using PBS) were maintained (shown in Fig. S2). The meniscus root-bone specimen was gripped with sandpaper. After preconditioning the specimen at 3% for 10 cycles using a strain rate at 0.1%/s, sample was loaded for 10000 cycles straining at 3% with loading rate of 0.5 Hz, which is the physiological strain amplitude ^13^. Slight decline observed on the relative load for >5000 cycles was possibly due to incomplete simulation of physiological environment by the chamber. Relative load plotted in results were obtained by normalizing real-time load to the load level during preconditioning.

### Fracture toughness estimation

Fresh porcine stifle joints were acquired from abattoir instantly after sacrifice and kept on ice until dissection. Meniscus-bone samples were dissected from the tibial plateau, with meniscus body removed within 12 hours. The meniscus root-bone samples were sectioned using a microtome blade to a thickness of 0.5-2 mm and incised into sizes of about 1 cm in length and width. Apart from the meniscus-bone interface, part of the root and bone segments were also included in the specimen for convenience of gripping samples. A cut at the interface with a width of 30-40% of sample width were made from the edge of the specimen before testing as the initial crack. Sandpaper were attached to grips for fixation of samples and renewed for each test. Samples were kept hydrated using gauze in PBS during testing. Representations of these configurations are shown in Fig. S29.

The meniscus-bone interface is a complicated composite composed of soft tissue (unmineralized) and hard tissue (mineralized). Over 40 samples were tested to estimate the fracture toughness of the interface. In observations, most specimens can resist crack propagation and development of crack only occurs in 10% of testing specimens, from which we collected crack propagation data. Since observable crack development occurred only in soft tissue region and given the fact that there is no standard fracture toughness test method for such a strong soft-hard interface, we applied the method for soft tissue’s fracture toughness to estimate the fracture toughness of the interface of interest(*64*).

The process for evaluating fracture toughness, denoted as *J_c_*, involves subjecting a test specimen with an initial notch to cycles of tensile loading and unloading, followed by dividing the energy required for crack propagation, *U_c_*, by the increment in crack length (δ*a*) multiplied by the thickness of the sample (*t*) to determine the newly formed crack area:

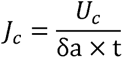

Ashby plot of fracture toughness and modulus data containing other materials were reproduced from published data(*29*).

### Interface adhesion test

Fresh porcine meniscus root-bone tissue specimens were obtained and sectioned into 2 mm thick pieces as described in the section of “fracture toughness estimation”. Specimens were segregated into 2 groups (n=5): one group underwent no treatment; the other group was segmented at the root-bone interface using a microtome blade, and the segmented halves were sealed back to the original state using cyanoacrylate. Being gripped through sandpaper, pull-to-failure tensile test was performed on both groups, with mechanical data and optical images of the testing process recorded.

### Atomic force microscope (AFM) imaging

The mechanical characteristics of the human meniscus root-bone interface were evaluated using atomic force microscopy (AFM) in QI mode with a NanoWizard 5 system from JPK Instruments AG, Berlin, Germany, immersed in PBS. Cryo-sectioned specimens with thickness of 10 μm were employed for AFM imaging. To investigate the samples, a paddle silicon nitride tip with a cantilever spring constant of approximately 0.1 N/m was utilized for unmineralized regions, while a pyramidal silicon nitride tip with a cantilever spring constant of around 42 N/m (TESPA-V2, Bruker) was employed for mineralized areas. Measurements were conducted by performing force-indentation ramps, with a 1 nN setpoint for soft regions and a 100 nN setpoint for stiff regions, employing a 30 x 30 µm scan range at a ramping speed of 2 µm/s. A grid comprising 128 x 128 pixels was created by executing a force-indentation curve for each pixel. Subsequently, the data were processed using JPK processing software to obtain parameters such as height, Young’s modulus, adhesive force, and dissipation energy.

### Scanning electron microscope (SEM)

50 μm cryo-sectioned specimens were thawed, washed with deionized water and dehydrated using gradient ethanol solution (30%, 50%, 70%, 80%, 90%, 100%-1, 100%-2). The specimens were incubated in each solution for 30 min and airdried. Glass slides were used to keep specimens flatten during airdrying. SEM (Hitachi, SU5000) was used to observe the specimens with the voltage of 10 kV after coating with platinum. In addition to Secondary Electrons (SE) mode, Backscattered Electron (BSE) mode was used to identify calcium-rich regions in organic matrix of the tissue.

### Element Dispersive Spectroscopy (EDS)

EDS spot scan, line scan and mapping scan were performed on selected positions in TEM. Ca/P ratio data was calculated from the atomic ratios of elements from spot scan data.

### Osmium tetroxide-thiocarbohydrazide-osmium (OTO) protocol of sample preparation

Meniscus root samples were trimmed to cubes with sides of 2 mm and underwent fixation in 2.5% glutaraldehyde for 24 h. 0.1 M PBS was used to wash the samples for three times. The samples were subsequently transferred to 1:1 mix solution of 2% osmium tetroxide and 3% potassium ferrocyanide and incubated for 1 hour. Samples were then incubated in 1% thiocarbohydrazide, 2% osmic acid and 1% uranyl acetate and Walton’s Lead Aspartate Solution with washing between each solution.

Dehydration was completed by gradient series of ethanol (30%, 50%, 70%, 90%, 100%-1, 100%-2) for 60 min in each solution. Dehydrated samples were washed in acetone which was then replaced with mixture of acetone and epon, and finally the samples were embedded in spur resin.

### High pressure freezing (HPF) protocol of sample preparation

The specimens were immersed in an external cryoprotectant solution (1-hexadecene) and gently placed into HPF sample carriers (depth: 0.2 mm, Type A, Leica 16770141). The cover (Type B, Leica 16770142) was also coated with 1-hexadecene and positioned over the sample carrier. Freezing was performed using an HPF apparatus (Leica EM ICE), followed by freeze substitution in EM AFS2 (Leica). Ethyl alcohol was used to fill the sample chamber to keep the temperature stable. The sample carrier was then immersed in 0.1% OsO4 solution in acetone. Following rinsing in acetone and substitution with a mixture of acetone and epoxy, the specimens were embedded in spur resin.

### Raman imaging and stimulated Raman scattering (SRS) microscopy

50-μm cryo-sectioned meniscus root-bone specimens were thawed, washed with PBS and transferred to clean slides and immersed in PBS for Raman spectroscopy imaging. 532 nm laser on Raman microscope (LabRAM HR Evolution, Horiba Co., Ltd.) was used to collect Raman spectra from 300 to 1800 cm^-1^ through the electron multiplying charge-coupled device (EMCCD) detector. Point data was obtained using a 5 s accumulation time whilst mapping data was acquired with 0.5 s accumulation time. Both pointing and mapping data had spatial resolution of 1 μm. We observed no destruction on tissue specimens under the working parameters mentioned above. SRS imaging was performed through Multimodal Nonlinear Optical Microscopy System (UltraView, Zhendian (Suzhou) Medical Technology Co., Ltd). Hyperspectral scan was utilized to obtain spectra at multiple ranges (800-1000 cm^-1^, 1300-1500 cm^-1^, 2800-3000 cm^-1^) from specimens prepared in the same way as Raman spectroscopy with a spectral resolution of 0.5 cm^-1^. Each scanning data was acquired in 400 x 400 pixels with spatial resolution of 0.52 μm/pixel and 30 μs dwell time. SRS intensity mapping of different constituents was acquired using the least absolute shrinkage and selection operator (LASSO) method as previously described(*65*). The Raman and SRS peak assignments are listed in Table S3.

### Focused Ion Beam (FIB)-SEM

The resin blocks underwent trimming on the Leica EM trimmer to expose the stained tissue surface. SEM imaging (Thermo Fisher, Teneo VS) was employed to identify the region of interest. An ultramicrotome located within the specimen chamber facilitated the milling of resin blocks and simultaneous acquisition of SEM images. Sample blocks were observed via FIB via a dual-beam scanning electron microscopy setup (Thermo Fisher, FIB Helios G3 UC). Serial images were acquired in the serial-surface view mode, with the thickness of slices set to 5 nm, at an energy of 30 keV and a current of 0.79 nA. Subsequently, each exposed serial surface underwent imaging at 2 kV (acceleration voltage) and 0.2 nA in backscattered electron scanning mode utilizing an ICD detector. The images were stored at a resolution of 3072×2048 pixels and the dwell time was 15 μs per pixel and pixel size set to 4.25 nm. Alignment and segmentation of the image stacks were executed using Amira 6.5 (Thermo).

### High-resolution Transmission electron microscope (HRTEM) and associated analyses

Sample blocks prepared according to the OTO protocol were sliced into 100-150nm thickness using Leica EM UC7 ultramicrotome. Sections were held on 100-mesh copper grids with a carbon support film. Sections were observed using an aberration-corrected scanning transmission electron microscope (Titan G2 80-200 microscope with a Super-X EDX detector, FEI) at 80 kV. High angle annular dark field-scanning TEM (HAADF-STEM), STEM-EDS mapping, Energy Dispersive X-ray Scanning (EDS) and selected area electron diffraction (SAED) patterns were combined with high-resolution TEM (HRTEM) during imaging. Electron energy loss spectra (EELS) and intensity images were collected at 100-600 eV to explore the featured edges of Phosphorus, Carbon, Calcium, Nitrogen, Oxygen. Calibration, normalization, background elimination and curve fitting were performed to analyze the EELS data (EELS characteristic peaks are listed in Table S4). The electron probe underwent calibration alongside a DCOR plus spherical aberration corrector, employing a standard sample for reference. SAED, HRTEM, EELS data were analyzed utilizing Gatan Digital Micrograph.

### Cryo-TEM imaging and 3D tomography

Sample blocks prepared according to the HPF protocol were sliced into 100 nm thickness using Leica EM UC7 ultramicrotome cryo-chamber maintaining the specimens under −150 °C. Cryo-TEM imaging and 3D tomography were conducted using Thermo Fisher Scientific Talos F200C at 200 kV in the cryo-mode. Tilt series for tomography were obtained at −60° to 60° with 1° steps and were subsequently aligned. Tomography data was further processed and visualized using Amira version 2019.1.

### LC-MS/MS Analysis

100-μm cryosections of meniscus root from three donors were washed with deionized water, and cut into three parts: bone, interface and root. Separated samples were transferred to 0.6-mL centrifuge tubes. 20 μl Ammonium bicarbonate (0.1 M) was added to each tube and heated at 95 °C for 10 min. Trypsin was added at 1 μg/μL for digestion at 37°C overnight. After centrifugation at 14000g for 15 min, the tryptic peptides in supernatant were quantified using NanoDrop Microvolume Spectrophotometers (Thermofisher). Peptide solutions were desalted using trifluoroacetic acid and C18-StageTip (Thermofisher) following manufacturer’s protocol. Desalted peptides underwent drying by vacuum centrifuging and were kept at −20°C upon analysis.

0.1% formic acid (solvent A) was utilized to dissolve the tryptic peptides. Subsequently, the peptides were introduced onto a reversed-phase analytical column filled with Reprosil-Pur C18 beads (1.9 μm, Dr. Maisch, Ammerbuch, Germany). A gradient elution strategy was employed using varying concentrations of solvent B (0.1% formic acid in 98% acetonitrile):starting from 3% to 10% over 3 minutes, increasing from 10% to 24% over 37 minutes, followed by a rise from 24% to 38% in 12 minutes, reaching 80% in 4 minutes, and maintaining at 80% for 4 minutes. Throughout, the flow rate was held constant at 450 nl/min using an UltiMate 3000 nanoLC system. The peptides underwent nanoelectrospray ionization (NSI) before tandem mass spectrometry (MS/MS) analysis on an Orbitrap Exploris 480 mass spectrometer (Thermo Fisher Scientific). The applying electrospray voltage was set to 2.0 kV. Full scan mass spectrometry was conducted over the m/z range of 400 to 1200 and the resolution was 60,000 in the Orbitrap. MS/MS fragmentation was conducted with a normalized collision energy (NCE) setting of 27, and fragment ions were identified in the Orbitrap using a resolution of 15,000. Data acquisition followed a data-dependent approach, changing between one MS scan and 22 MS/MS scans, with a dynamic exclusion of 30 seconds. The automatic gain control (AGC) was 5E4, and the compensation voltage for FAIMS was adjusted to −45V.

### Protein identification and quantification of LC-MS/MS data

Utilizing the MaxQuant search engine (version 1.6.15.0), the acquired MS/MS data underwent processing. Tandem mass spectra were compared against the Uniprot Human database and a reversed decoy database. The cleavage enzyme Trypsin/P was assigned, permitting up to 2 missed cleavages. For the first search, the tolerance for precursor ion mass was set at 20 ppm, while for the Main search, it was set to 5 ppm. Fragment ion mass tolerance was established at 0.02 Da. Fixed modification included Carbamidomethyl on cysteine, while oxidation on methionine was treated as a variable adjustment. Quantification without labelling was executed using the LFQ method, with a false discovery rate (FDR) fine-tuned to < 1%, and a least peptide score threshold set at > 40. The resulting data in the proteingroups.txt file were analyzed using the DEP package in R Studio. Three biological replicates were utilized for analysis. Any potentially contaminated samples, reverse data, and duplicated gene names were removed. Rows of proteins were filtered to retain only those with at least two out of three valid values found in individuals per group. Data normalization was carried out using variance stabilizing transformation followed by log2 transformation. K-nearest neighbors’ algorithm was applied to impute missing values. Linear Models and Empirical Bayes methods were used to obtain the differential expression of proteins (fold change >2, adjusted p value < 0.05). Proteins enriched in interface more than twofold were categorized by searching the protein name in genecards.org.

### Immunofluorescent staining

15-μm cryosections of samples were fixed with 4% paraforalderhyde for 10 min and washed with PBS. After incubating with 1% bovine serum albumin (BSA) in PBS, the samples were treated with primary antibodies including MGP (1:400, 60055-a-Ig, Proteintech, USA), HTRA1 (1:50, 55011-1-AP, Proteintech, USA) diluted with 0.1% BSA at 4 °C overnight. Followed by PBS wash, samples were stained with secondary antibodies (Goat Anti-Mouse Alexa Fluor® 647, ab150115; Goat Anti-Rabbit Alexa Fluor® 555, ab150078) at room temperature for 60 min, and a subsequent incubation with 0.01 μg/ml calcein (Solarbio, China) solution for 120 min. After mounting, sections were observed via a confocal microscope (Zeiss 880, USA).

### In vitro mineralization

Nanosized ACP clusters (nano-ACP) was prepared according to reported methods(*66*). 200-mesh golden grids with a carbon support film were added with 3 μL recombinant matrix gla protein (MGP) (100 μg/mL, CUSABIO, China) diluted with deionized water and the control group was added with 3 μL deionized water, and both were incubated for 20 minutes at room temperature. The grids were subsequently overturned and floated on nano-ACP solution and incubated at 37 °C. Grids from both groups were collected after 24 hours and 72 hours and the changes were observed through TEM (FEI Tecnai G2 F20 S-TWIN, Thermo) operating at 80 kV. Mineral density was quantified by measuring dark areas in TEM images for each group (n=5) using ImageJ and performing statistical analysis (student’s t test).

### *In situ* tensile test under fluorescent microscope

30-μm thick human meniscus root-bone tissue sections were thawed and washed with PBS. Then the samples were labelled with sulfo-Cyanine3 NHS ester (1:1000, Duofluor, China) or Calcein (1:500, Solarbio, China) for whole tissue and mineralized tissue visualization respectively. Intact stained sections at S-H interface were exposed to tensile stress in PBS bath under fluorescent microscope through a customized chamber set up on fluorescent microscope Nikon Eclipse Ni-E (Nikon, USA) as illustrated in Fig.S20. Stepping motor (PDV, China) and mechanical sensor EVT-FT200A (Shanghai Yu Ran Sensor Technology CO., LTD., China) were equipped in the chamber. Sample sections were preconditioned the at 3% for 5 cycles using a strain rate at 0.1 mm/s. Tensile stress was loaded on the section at 0.1 mm/s and tissue deformation was recorded in the form of videos. Screenshots from the videos were processed and displacement field was analyzed by PIV algorithm in ImageJ(*8*) using parameters shown in Table s5. Lagrangian strain field was analyzed and visualized by ncorr in matlab.

Sample sections with pre-created cracks and underwent tensile loading until failure were fixed immediately after loading using 4% PFA for 15 minutes and washed with deionized water. Subsequently the sections were dehydrated and observed in SEM as described in the “SEM” section.

### *In situ* tensile test in SEM

Human meniscus root-bone tissue pieces with a thickness of 2-4 mm were trimmed to dogbone shape, freeze-dried and coated with platinum. The samples were then transferred to in-situ tensile test system (Qiyue tech. (Hangzhou)) in Field Emission Scanning Electron Microscope (TESCAN CLARA). Samples were strained to 5%, 10%, 15% …at 1 μm/s until failure. Straining was paused per 5% strain to take snapshots at locations of concern (mineral particles in partially mineralized region of S-H interface) at higher magnification. The force-displacement curves, snapshots and videos of the whole process were obtained simultaneously.

### Molecular dynamics modelling

#### 1. Construction of collagen molecular model

Collagen’s full atomistic structures were assembled utilizing the sequence data from Pubmed. I-TASSER server was employed to align the sequences with crystallography-derived collagen structures, selecting the structure with the lowest energy. The resulting triple helix had a diameter of 1.28 nm and a length of 298.025 nm.

#### 2. Construction of ACP molecular model

The formula of ACP was set as Ca_2_(HPO_4_)_3_^2–^, referring to sphere-shaped aggregates reported in a former observational study of ACP transforming into apatite in biomimetic environment by Habraken et al.(*51*). This proposed formula was utilized due to two reasons: (1) the sphere morphology observed in the former study had strong resemblance with ACP particles found in meniscus root-bone interface in this study; (2) the formula was calculated through real-time monitoring of chemical changes in crystallization process, which render the results more analogous to the biomineralization process.

#### 3. Modelling of ACP and collagen

Molecular dynamics (MD) simulations were conducted utilizing the Gromacs 2019 version and energy of the system was minimized. Subsequently, the entire system was subjected to minimization while applying positional restraints to the backbone atoms of collagen. Following this, equilibration was conducted at 310 K for 500 picoseconds under constant number, volume, and temperature (NVT) conditions, followed by an additional 500 picoseconds under constant number, pressure, and temperature (NPT) conditions (temperature = 310 K, pressure = 1 atm). Throughout these equilibration phases, a positional restraint of 2 kcal mol-1 Å-2 was applied to the main chain carbon atoms of PAA to facilitate relaxation of its side chains. Finally, a 10 nanosecond MD simulation was performed on the systems in the NPT ensemble (temperature = 310 K, pressure = 1 atm) with periodic boundary conditions. To observe the interactions between ACP and collagen, dynamic changes in the ACP around the collagen structures were plotted (Fig.5g).

### Finite element analysis (FEA)

#### 1. FEA of meniscus root-bone tissue

Based on AFM modulus data, a simplified plane-stress model was constructed in ABAQUS to elucidate the strain and stress distributions within the root-bone interface under tensile loading (Fig. 1J). The model had a length of 180 μm and a width of 15 μm, containing tissue modulus transition from ligamentous root to mineralized fibrocartilage. The AFM data covers consecutive regions mapped from the fibrocartilage to the mineralized fibrocartilage, while the ligamentous root region is located ~ 200 μm away from the fibrocartilage region (demonstrated in Fig.S5). Therefore, a transitional zone was generated between the ligamentous root region and fibrocartilage region. Axial displacement equivalent to a total strain of 10% was applied on the boundaries and the maximal principal strain distribution was shown in Fig. S4 and Movie S1. From the results and modulus distribution, deformation mainly happens within the ligamentous root region which possesses low modulus, while the stiff mineralized region remains significantly undeformed.

#### 2. FEA of FIB-derived collagen and mineral microstructure

Finite element analysis (FEA) with ABAQUS simulated the uniaxial stretching of collagen-mineral structures. A 3D model, representing the mineral-collagen fibril composite, was derived from FIB-SEM data of the meniscus root’s S-H interface and imported into ABAQUS. Tetrahedral elements meshed the 3D structures, using Amira Meshing (2019.1, Thermo), ensuring no overlap between mineral particles and fibrils in extrafibrillar space. Mineral particles exhibited a linear elastic behavior with a Young’s modulus of 125 GPa and a Poisson’s ratio of 0.3, while collagen fibrils had a mean modulus of 2.5 GPa with a Poisson’s ratio of 0.4.

The finite element models (FEMs) utilized four-node plane-strain quadrilateral elements (CPE4) from the ABAQUS library. We refined the mesh around defects for precise localized effect representation and used a coarser mesh elsewhere. Elements varied from a quarter to 20 units horizontally and one unit vertically. Strain loads of 1%, 3%, 5%, 7%, and 9% were applied along the axial direction of fibrils, with displacement constraints imposed on the lateral boundaries orthogonal to the load direction. The resulting stresses and strains were visualized on the 3D mesh, and data from 20 random points in contact and non-contact areas were extracted for further analysis.

### Over-mineralization and demineralization of root-bone interface

Porcine root-bone interface tissue specimens were harvested from fresh porcine stifle joints as described in ‘Fracture toughness estimation’ section. After washing with PBS, root-bone specimens were immersed into demineralization reagent 10% Ethylenediamine tetra-acetic acid disodium salt dihydrate (EDTA, Sinopharm Chemical Reagent Co., Ltd.) or over-mineralization reagent. The over-mineralization reagent was prepared as follows: in a culture dish, mix 1 mL of 3.34 mM [Ca] solution (from CaCl2) with 1 mL of 19 mM [P] solution (a mixture of Na_2_HPO_4_ (Sigma) and 300 mM NaCl (Sigma)). Add 48 μL of 10 mg/mL p-Asp to reach a final concentration of 240 μg/mL. Filter both [Ca] and [P] solutions through a 0.22 μm membrane and mix the cold solutions by adding [Ca] first, then p-Asp, and finally [P]. Samples (n=6) in both demineralization and over-mineralization reagent were incubated at 37°C and the solution level were maintained even with the interface (over mineralized tissue). Control samples were immersed in PBS. Reagents were refreshed every 24 hours. After 48 hours, sizes of the samples were recorded and the samples were used for pull-to-failure test for quantification of strength and fracture toughness test for quantifying toughness.

### Mineralized artificial root fabrication

#### 1. SilMA preparation

50 g raw silk (Bombyx mori, Zhejiang Xingyue Biotechnology Co., Ltd., China) was boiled at 100 °C for 30 minutes in 0.05M Na_2_CO_3_ (Sigma) solution and this procedure was performed 3 times to eliminate the sericin. The degummed silk fiber was dissolved in 9.3 M Lithium bromide (Macklin) at 60 °C for 240 min with rotational rate of 300 rpm. Subsequently, the mixture received an infusion of 30 mL of glycidyl methacrylate solution, administered gradually at a pace of 0.5 mL per minute. The solution was subjected to filtration dialysis against distilled water using 12 kDa cutoff dialysis tubes for 4 days during which water was changed 3 times per day. The solution was then frozen at −80°C for 4 h, freeze-dried for 4 days and stored at 4 °C. Finally, the 15% silMA (w/v) solution was obtained by dissolving the freeze-dried silk fibroin into water for further use.

#### 2. Artificial ligament fabrication

The aligned silMA scaffolds were fabricated using directional freezing by injecting 650 ml 15% silMA solution into cylindrical plastic tube (5 mm inner diameter and 3 cm length), the bottom of which was in contacted with the copper rod pre-cooled using liquid nitrogen. The scaffolds were exposed to UV light for initial gelation and subsequently immersed in 100% ethanol for 24 h and in deionized H_2_O for 24 h to remove residual ethanol. For combined ACP and HAP mineralization, the bottom 1/3 portion (about 6 mm) was immersed in 10% (w/w) HAP (Aladdin) solution for 24 h, and then transferred to 5% (w/w) ACP (Sigma) solution which volume was over 2/3 height of the scaffold (about 12 mm) for 24 h, leaving the upper 1/3 region blank. Blank control scaffolds were immersed in deionized water for 48h. ACP-mineralized scaffolds and HAP mineralized scaffolds were immersed in 5% (w/w) ACP or 10% (w/w) HAP solution for 48 h. After mineralization, tensile tests were performed and tensile modulus and toughness were derived from stress-strain curves.

### Statistical analysis

All data were presented as the mean ± SD. Statistical differences among multiple (>2) groups were examined using one-way ANOVA analysis (Tukey’s posthoc test) and Students’ t test was utilized to compare data between two groups. p value < 0.05 was considered statistically significant.

## Code availability statement

The ImageJ PIV plugin can be accessed through the website: https://sites.google.com/site/qingzongtseng/piv. Ncorr algorithm and guidance are available at https://ncorr.com/index.php.

## Data availability statement

All data generated or analyzed during this study are included in this published article (and supplementary information files).

## Acknowledgements

We thank Nianhang Rong for the SEM imaging and EDX analysis. We thank Guoqing Zhu and Qinyun Lin from the Centre of Electron Microscopy, Zhejiang University for HRTEM, SAED and EELS imaging. We thank Jiansheng Guo and Lingyun Wu from the Centre of Cryo-Electron Microscopy (CCEM), Zhejiang university for technical assistance on Sample preparation, FIB-SEM imaging and cryo-TEM imaging. We thank Vibronix (Suzhou) for support in Stimulated Raman Scattering imaging. We thank Bruker (Shanghai) for providing support in AFM imaging. We thank Bin Gao for assistance in analysis of proteomics data. The authors also thank Pengda Zou and Minghui Li (Mass Spectrometry Core Facilities, The First Affiliated Hospital, Zhejiang University School of Medicine) for their assistance with LC-MS/MS. Funding from National Key Research and Development Program of China grant 2023YFB3813000 (H.O.), National Natural Sciences Foundation of China grant T2121004, 82394441, 92268203 (H.O.), 32371411 (X.W.), Key Research and Development Program of Zhejiang 2024SSYS0026 (H.O.) and AI Singapore grant AISG2-GC-2023-010 (DL) is gratefully acknowledged.

## Author contributions

W.L., X.W., H.O., and W.C. contributed to conceptualization. W.L., X.W., R.M., D.L., H.M., Z.K., B.W., W.J., R.L., and X.P. contributed to methodology. W.L., X.W., R.M., D.L., and H.M. contributed to investigation. W.L., X.W., R.M., D.L., H.M., R.Z., and C.X. contributed to visualization. J.F., W.D., and X.C. contributed to resources. R.M., D.L., R.Z., Z.K., and W.C. contributed to software. W.D. and H.G. contributed to supervision. W.L., X.W., and X.Y. contributed to writing and reviewing.

## Additional information

Supplementary Information is available for this paper.

Correspondence and requests for materials should be addressed to Hongwei Ouyang.

