## Supplementary Figures & Tables for "A Tough Biointerface in Human Knee Empowered by Dynamic Phase-transforming Minerals in Collagenous Matrix"

**The PDF file includes:**

Figs. S1 to S29

Tables S1 to S5

References

**Other Supplementary Materials include the following:**

Movies S1 to S3

Data S1

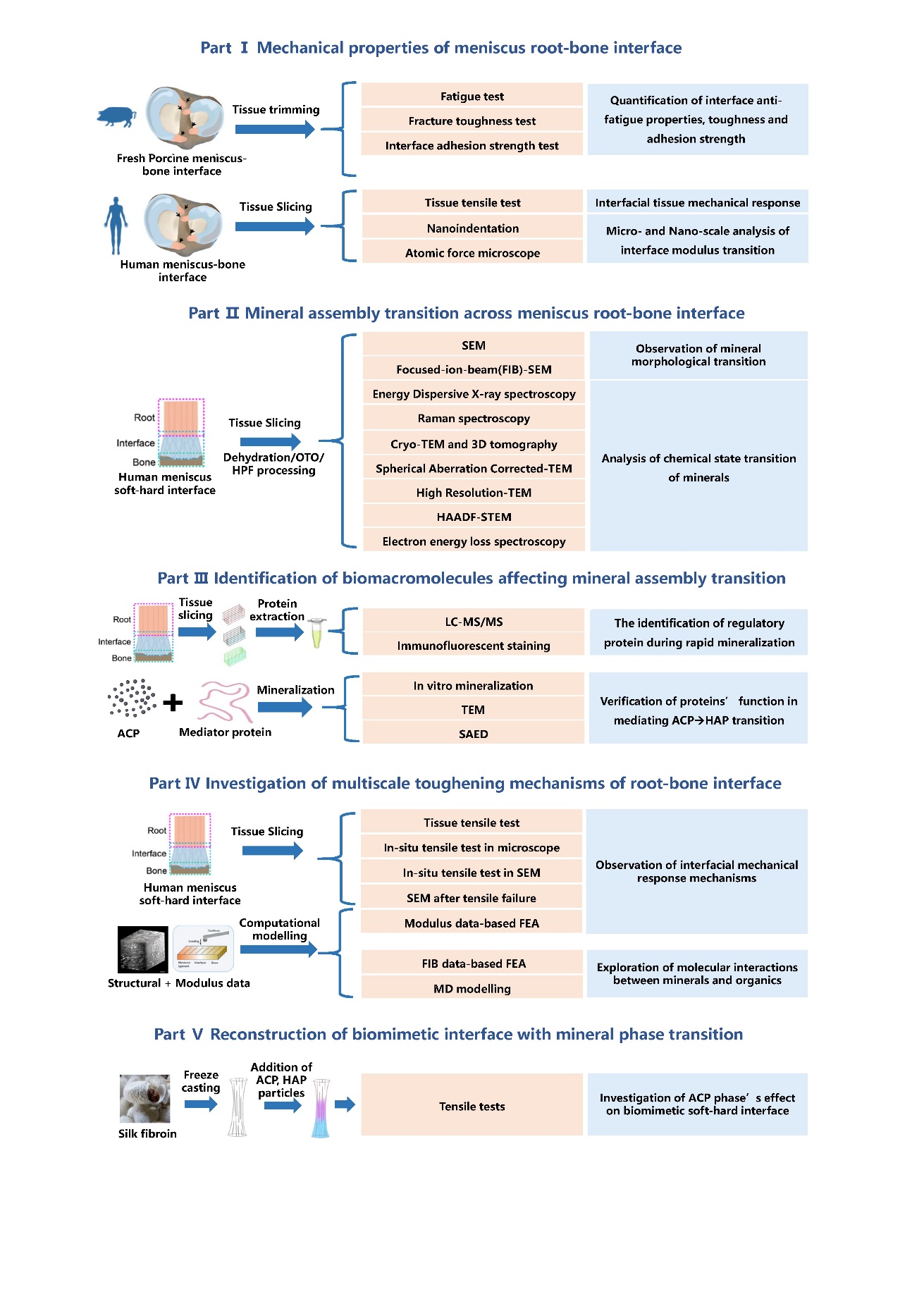

**Fig. S1**. Flow diagram illustrating the experimental procedure followed in this study. Abbreviations: OTO, Osmium tetroxide-thiocarbohydrazide-osmium method; HPF, high-pressure freezing; Cryo-TEM, cryogenic transmission electron microscopy; HAADF-STEM, high-angle annular dark field scanning transmission electron microscopy; LC-MS/MS, liquid chromatography−tandem mass spectrometry; SAED, selected area electron diffraction; SEM, scanning electron microscopy; FEA, finite element analysis; MD, molecular dynamics; ACP, amorphous calcium phosphate; HAP, hydroxyapatite.

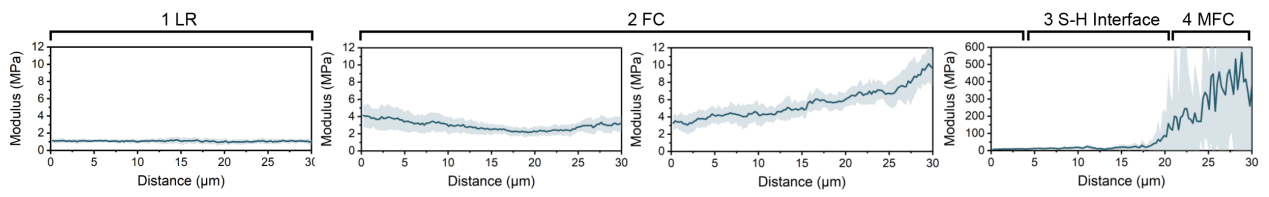

**Fig. S2**. Tissue modulus transition across the root-bone interface from ligamentous root (LR), throught fibrocartilage (FC) and soft-hard (S-H) interface, to mineralized fibrocartilage (MFC).

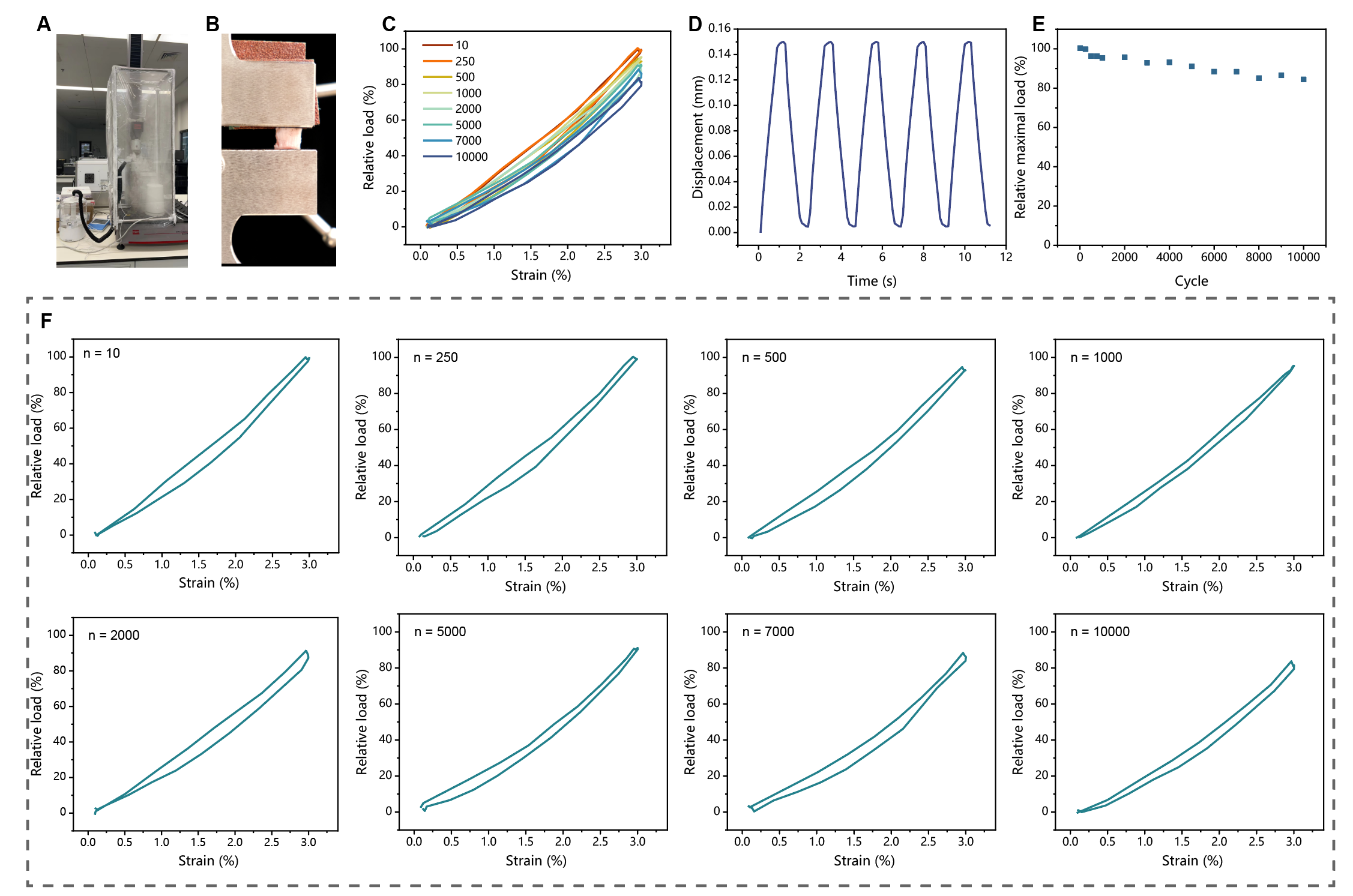

**Fig. S3**. Fatigue tensile test of the meniscus root-bone interface tissue. (**A**) Optical photo of the constant temperature and humidity chamber equipped on the loading apparatus. (**B**) Optical photo of the root-bone tissue clamped for fatigue loading. (**C**) Loading curves for various cycle numbers. (**D**) Cyclic loading and unloading curves showing loading frequency. (**E**) Relative load maintained at a steady level throughout cyclic loading. (**F**) Curves of relative load and displacement for a specified cycle number (*n*).

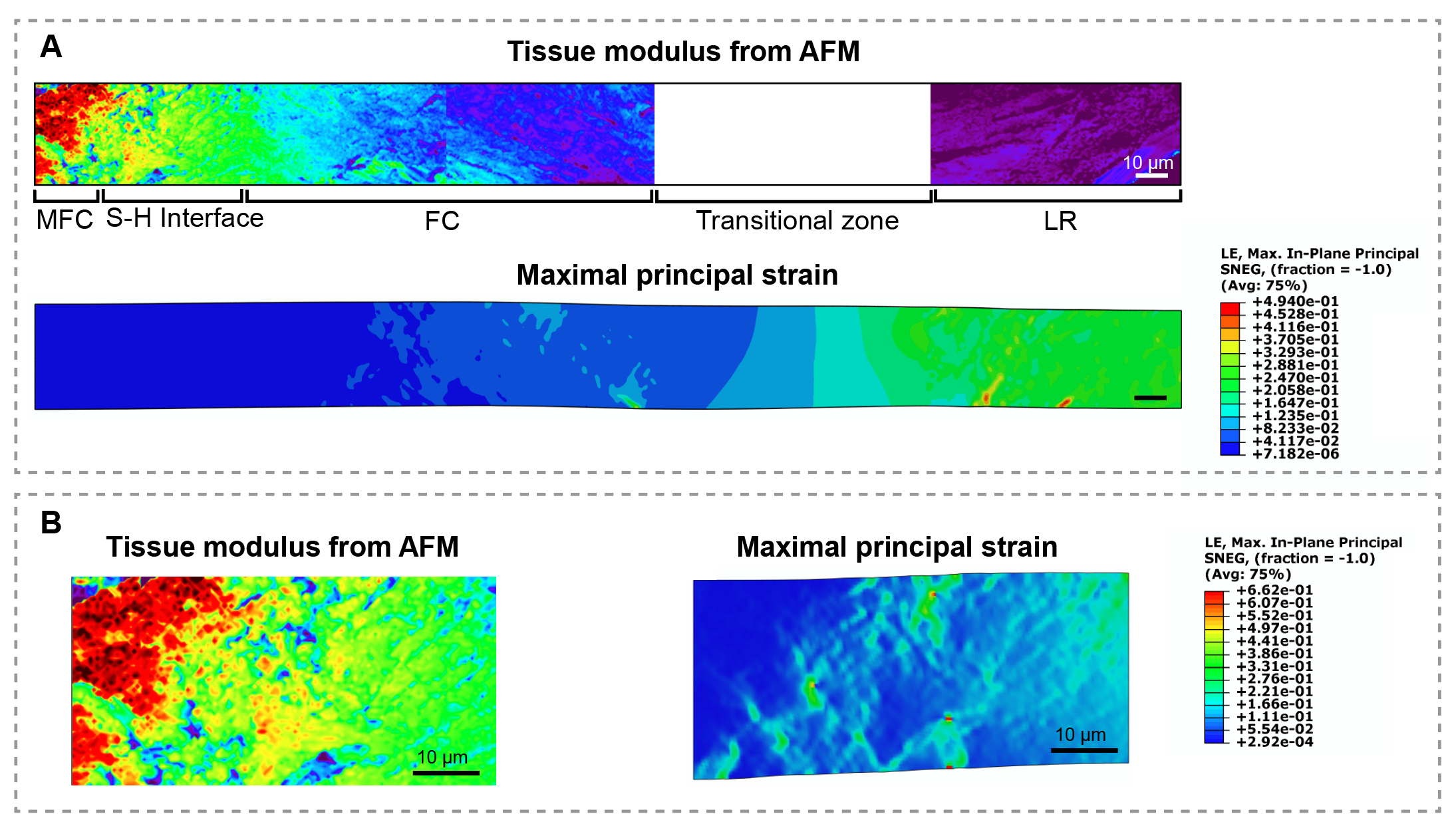

**Fig. S4**. FEA of the root-bone interface two-dimensional model under tensile stress. (**A**) The two-dimensional model was established based on AFM modulus data of root-bone tissue from the ligamentous root (LR), through fibrocartilage (FC), to mineralized fibrocartilage (MFC). A gap existed between the LR and FC regions, so a transitional zone with a smooth modulus transition was set, joining the FC and LR region in the model. (**B**) Enlarged view of the modulus and strain of the S-H interface region, showing that strain was concentrated in the unmineralized tissue.

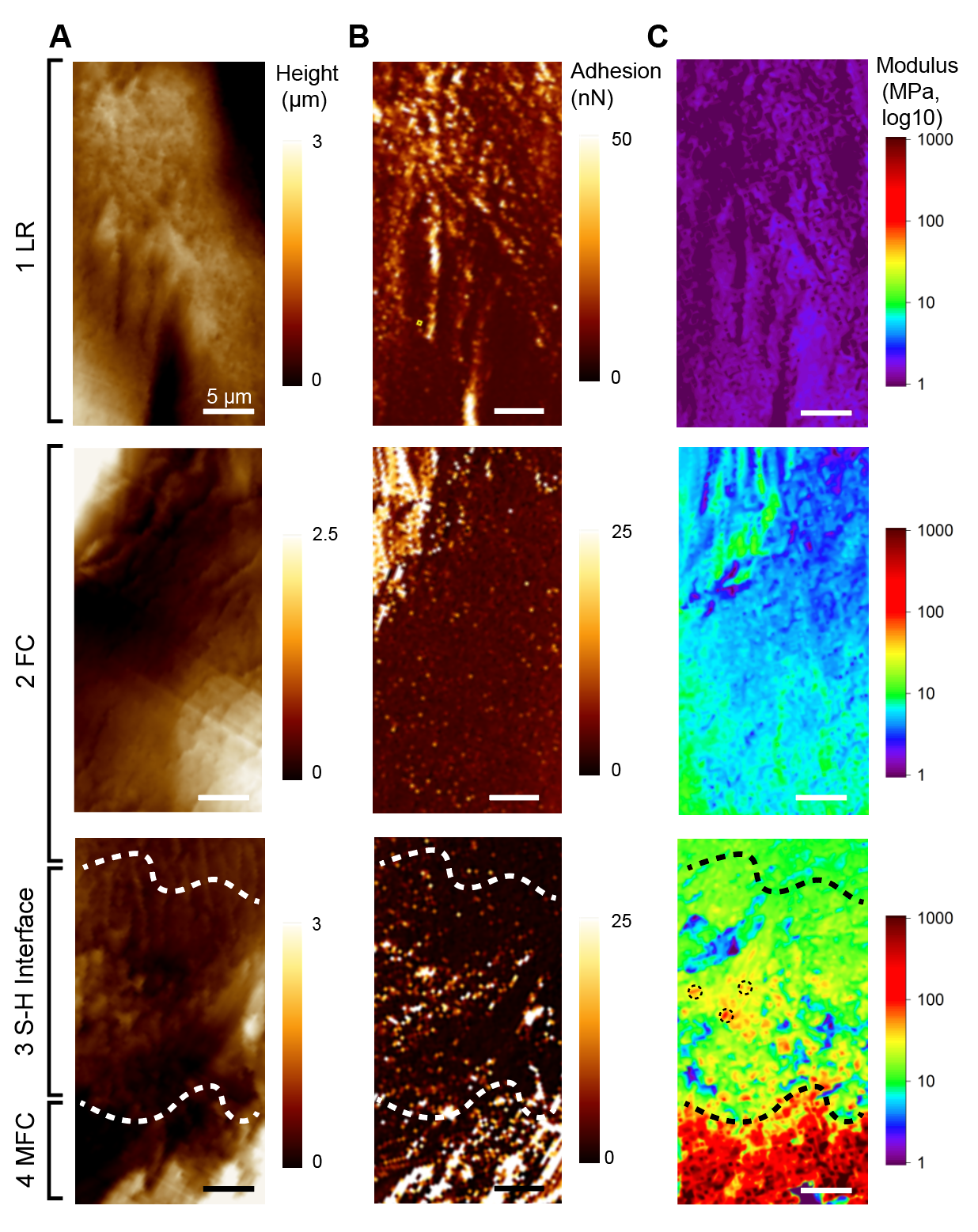

**Fig. S5**. AFM map of the root-bone interface tissue. Tissue height (**A**), adhesion strength (**B**), and modulus (**C**) maps of the ligamentous root (LR), fibrocartilage (FC), soft-hard (S-H) interface and mineralized fibrocartilage (MFC). White dashed lines indicate the S-H interface region, where the modulus gradually increased in a granular manner (indicated by black circles).

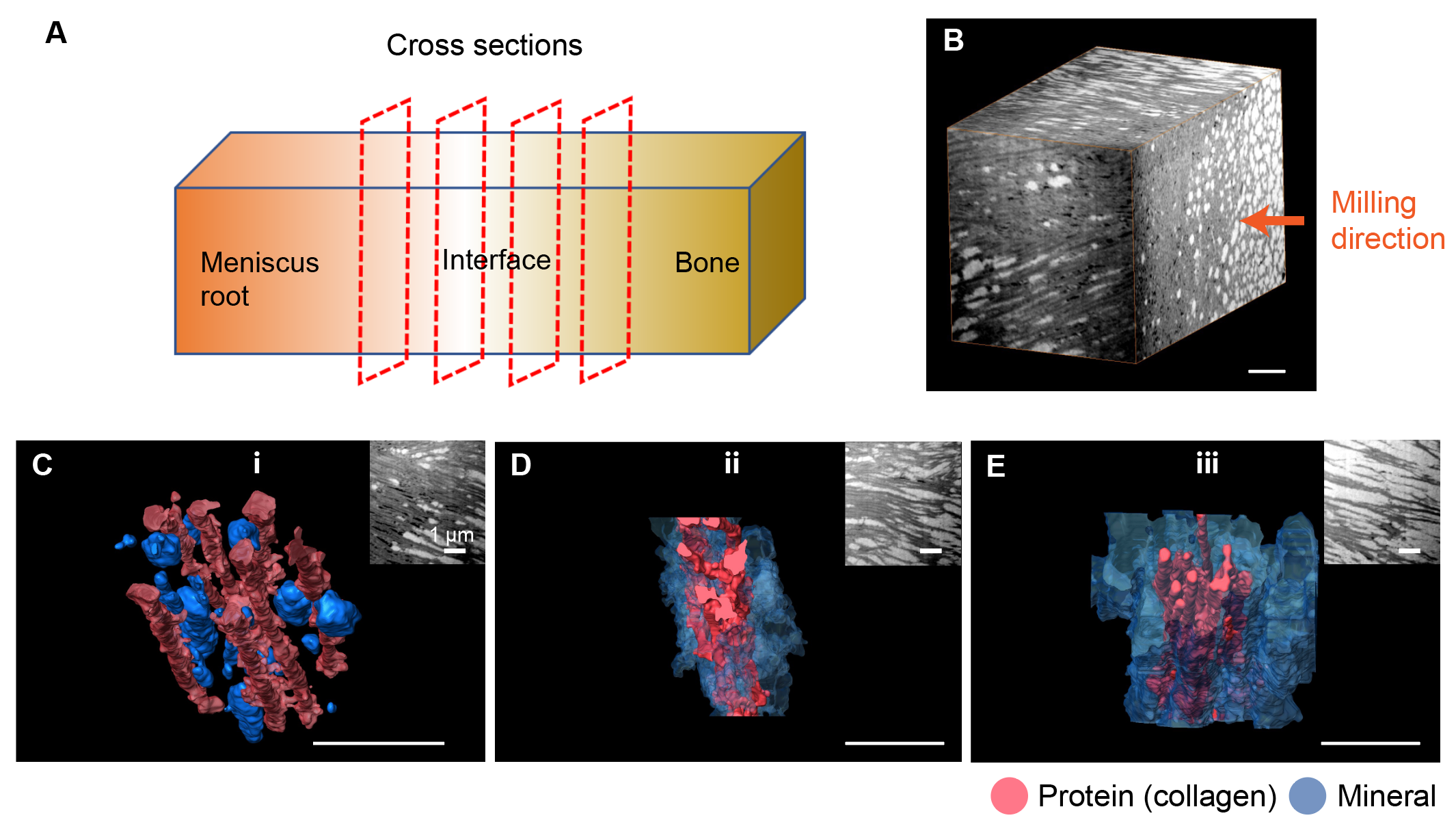

**Fig. S6**. FIB data of the S-H interface collected in the cross-sectional direction. (**A**) Schematic diagram showing the direction of the FIB images. (**B**) 3D view of the dataset with the FIB milling direction indicated by an orange arrow. (**C-E**) 3D reconstruction of region i-iii in this dataset, demonstrating the morphological and distributional transition of minerals. Inlets show the 2D view of the corresponding reconstructed 3D models.

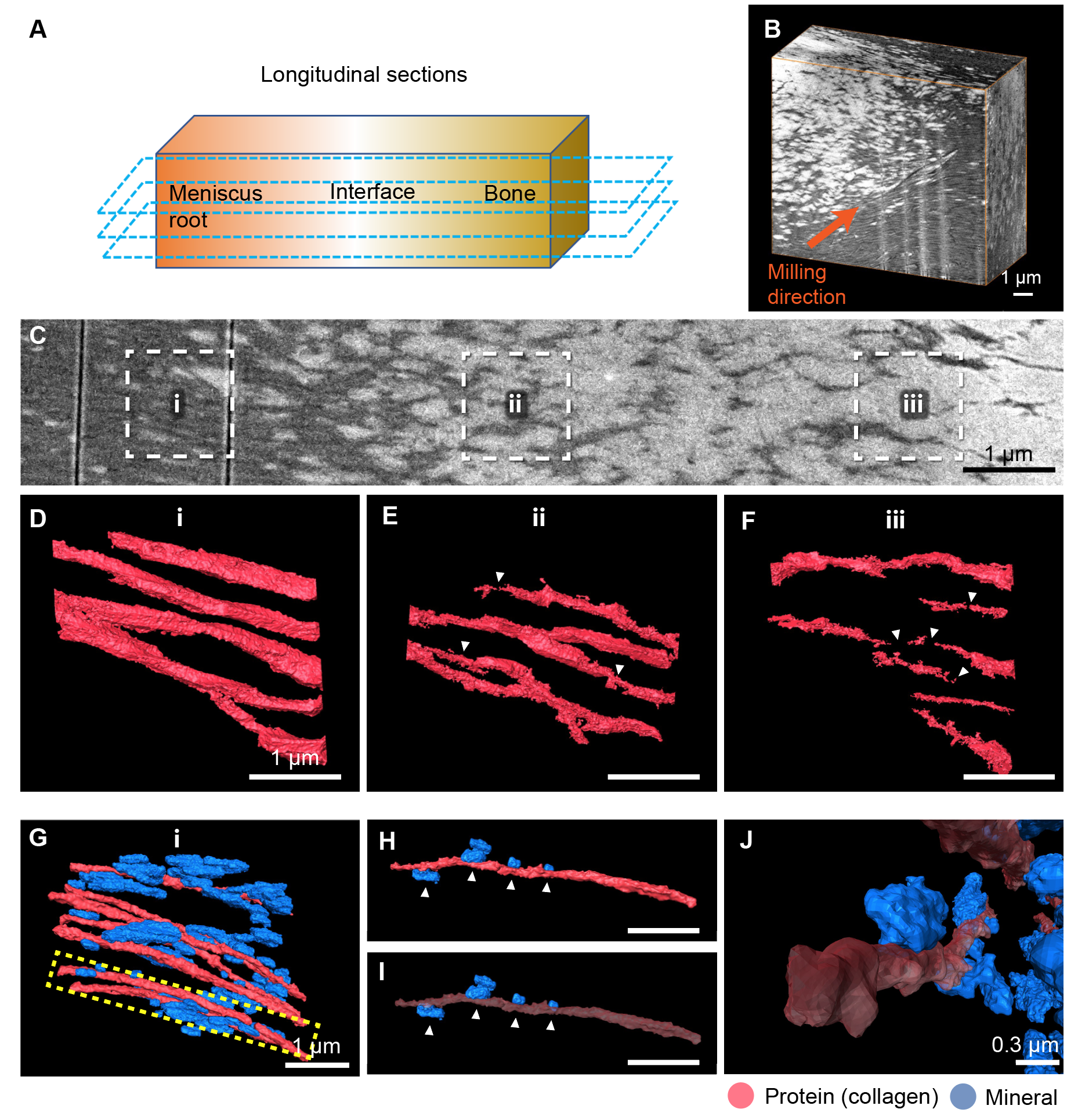

**Fig. S7**. FIB data of the S-H interface collected in the longitudinal direction. (**A**) Schematic diagram showing the direction of the FIB images. (**B**) 3D view of the dataset with the FIB milling direction indicated by an orange arrow. (**C**) 2D image of minerals with increasing density in region i-iii across the S-H interface. (**D-F**) 3D reconstruction of representative collagen fibrils in region i, ii and iii, respectively. Triangles denote partial deficiency of fibril structure, and this phenomenon is intensified from region i to iii, as minerals progressively occupy the intrafibrillar space. (**G**) 3D reconstruction of presentative structures in region i, revealing the relative location of mineral and fibrils. One of the fibrils is separately presented in (**H**) and (**I**), with the fibril set to 50% transparent in (i), showing minerals located in the periphery along the fibril. (**J**) Top view of the fibrils and minerals validates that mineralization only occurs outside fibrils in this region.

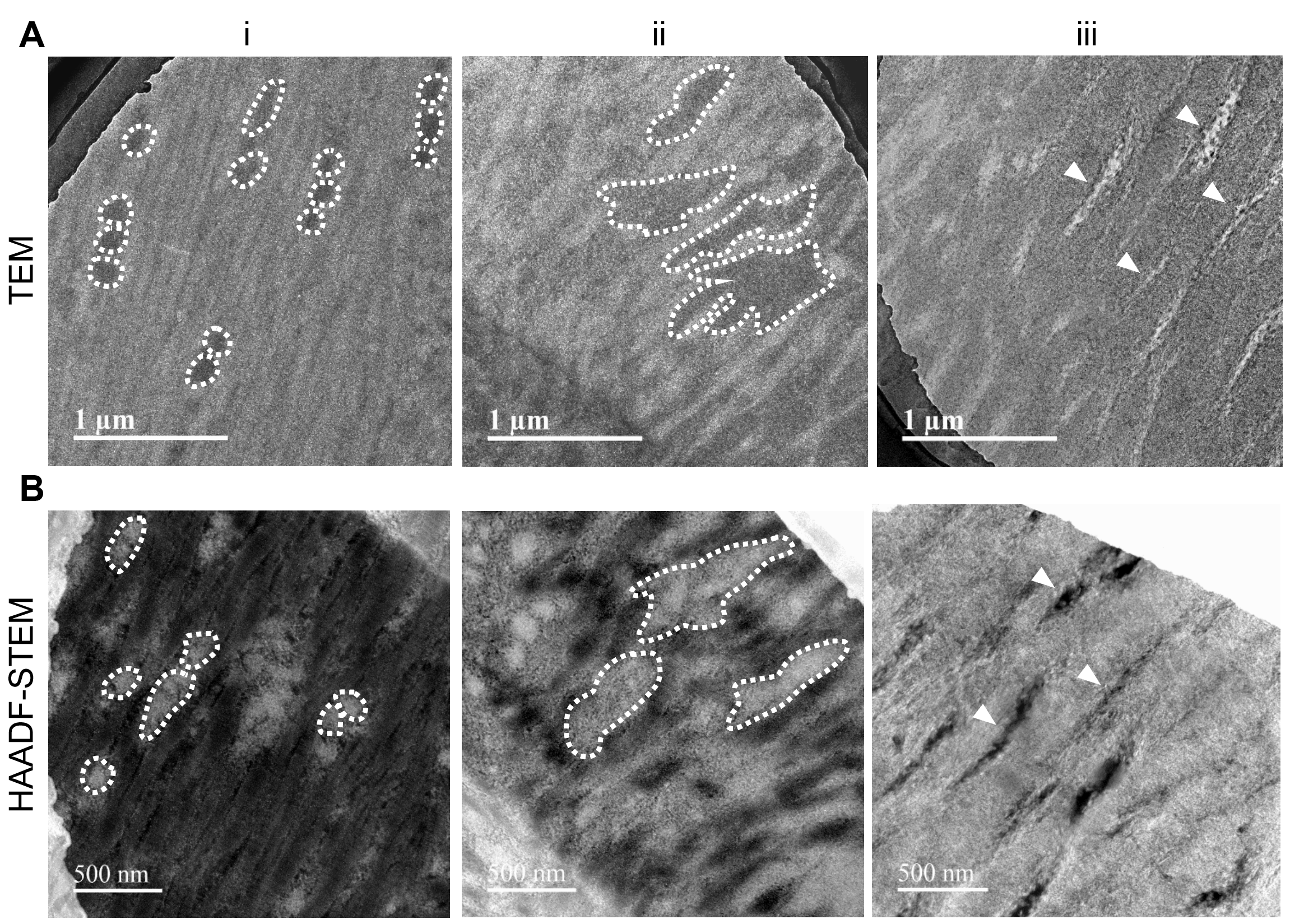

**Fig. S8**. TEM and HAADF-STEM images of minerals at the S-H interface. Minerals containing Ca and P were shown in relatively darker areas in (**A**) TEM and brighter areas in (**B**) STEM. Spherical mineral particles are indicated in dashed line in region i. Fused, irregular-shaped minerals can be found in region ii (also indicated by dashed line). In region iii, minerals occupy most of the space, except for a few gaps in the shape of deficient collagen fibrils (indicated by triangles). Abbreviations: TEM, transmission electron microscopy; HAADF-STEM, high-angle annular dark field scanning transmission electron microscopy.

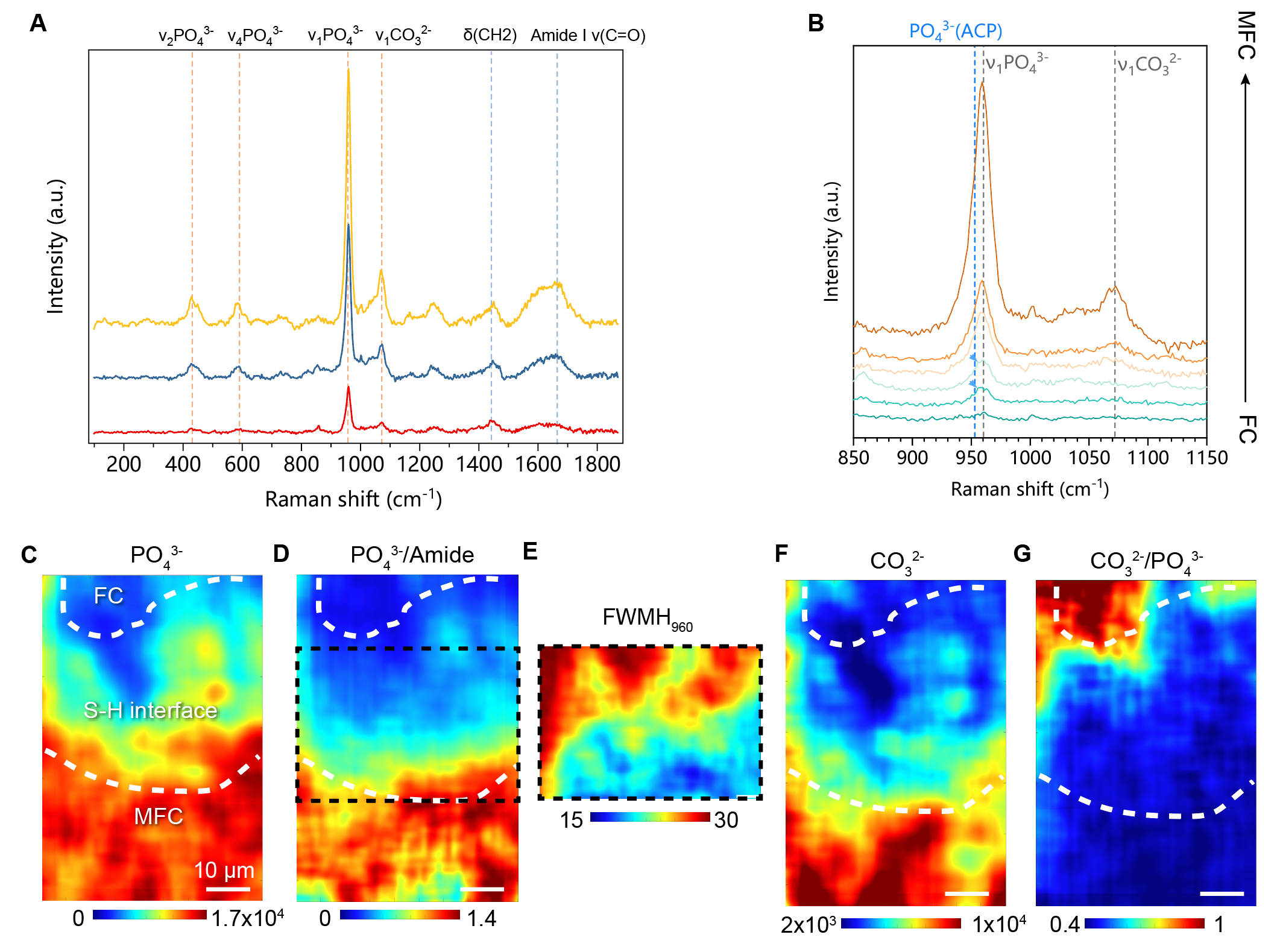

**Fig. S9**. Raman spectroscopic imaging of the S-H interface revealing compositional gradients from FC to MFC. (**A**) Overview of representative Raman peaks. ν_2_PO_4_^3-^, ν_4_PO_4_^3-^, ν_1_PO_4_^3-^ and ν_1_CO_3_^2-^ indicates the content of HAP and carbonate group, while δ_(CH2)_ and Amide I peak assignments indicate collagen protein. (**B**) An enlarged view showing shift from 960 cm-1 to the ACP peak (945-952 cm^-1^) observed at lower peaks of ν_1_PO_4_^3-^ where the mineralization degree is minor. Raman imaging of mineral content (**C**) relative to proteins (**D**) illustrates a gradient increase in mineralization degree. (**E**) Full-width of maximum height (FWMH) indicates the crystallinity of HAP. A decrease in FWMH from FC to MFC suggests improved crystallinity. (**F-G**) Carbonate content and carbonate substitution of HAP, which have a characteristic distribution over the interface. Abbreviations: MFC, mineralized fibrocartilage; FC, fibrocartilage; S-H interface, soft-hard interface.

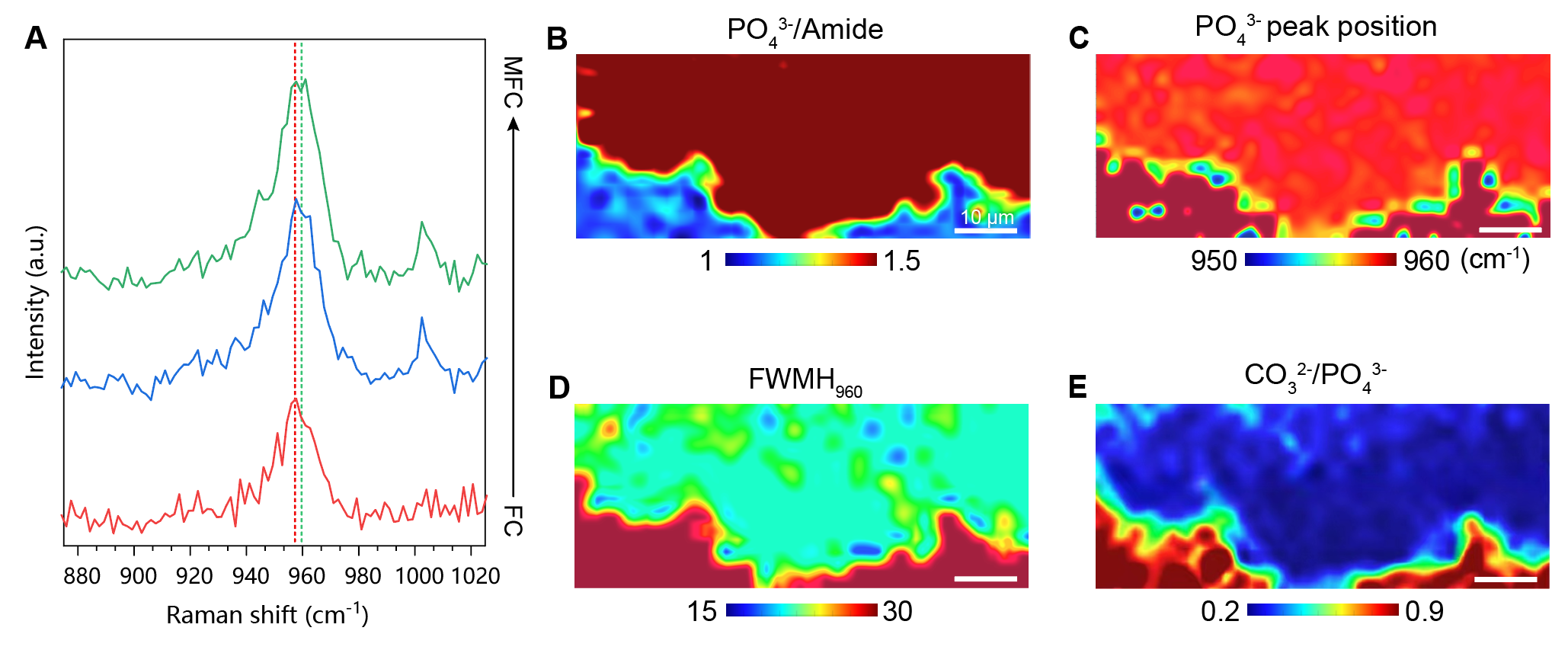

**Fig. S10**. Raman spectroscopic imaging of S-H interface from FC to MFC in porcine root-bone samples. (**A**) Representative Raman signals from FC to MFC regions showing shift from 960 cm-1 to the ACP peak (945-952 cm^-1^) where the mineralization degree is minor. (**B**) Raman imaging of mineral content relative to proteins. (**C**) Imaging of ν_1_PO_4_^3-^ peak position. (**D**) Full-width of maximum height (FWMH) indicates the crystallinity of HAP. (**E**) Imaging showing carbonate substitution of HAP.

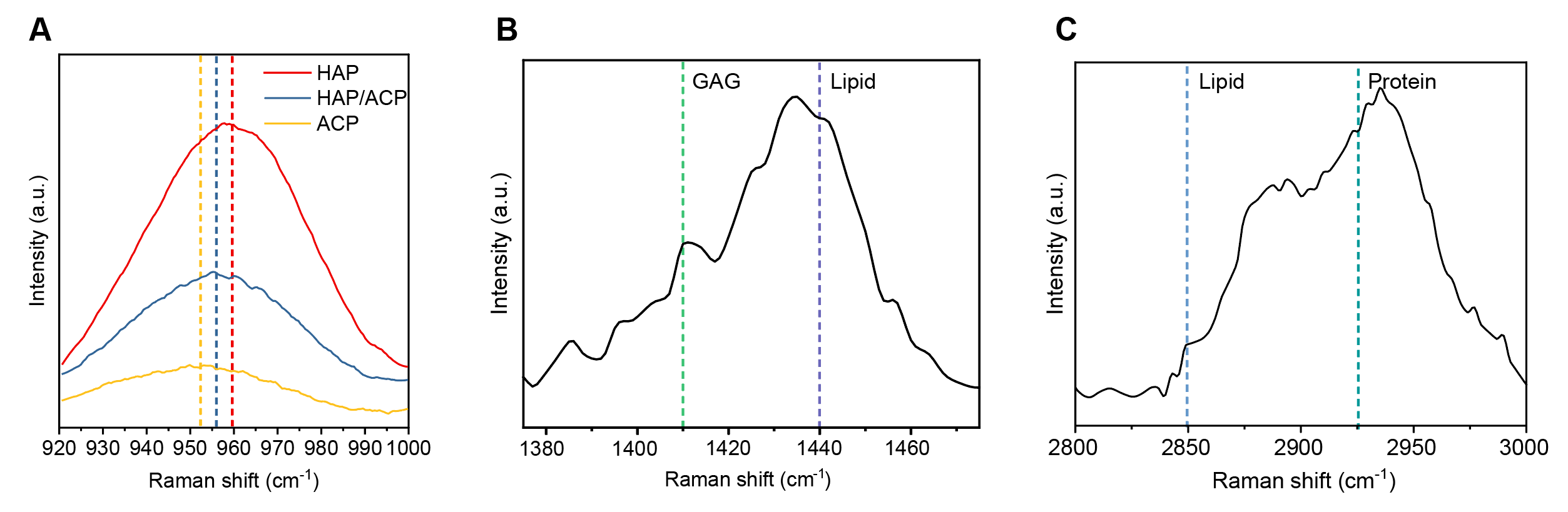

**Fig. S11.** Representative SRS curves used for intensity mapping. (**A**) ν_1_PO_4_^3-^ signals of ACP (952 cm^-1^), HAP (960cm^-1^) and their mixture (957 cm^-1^). (**B**) Peaks of GAG at 1410 cm^-1^ and lipid at 1440 cm^-1^. (**C**) Spectra of lipid and protein at 2850 cm^-1^ and 2926 cm^-1^ respectively. Abbreviations: SRS, Stimulated Raman Scattering; GAG, glycosaminoglycan.

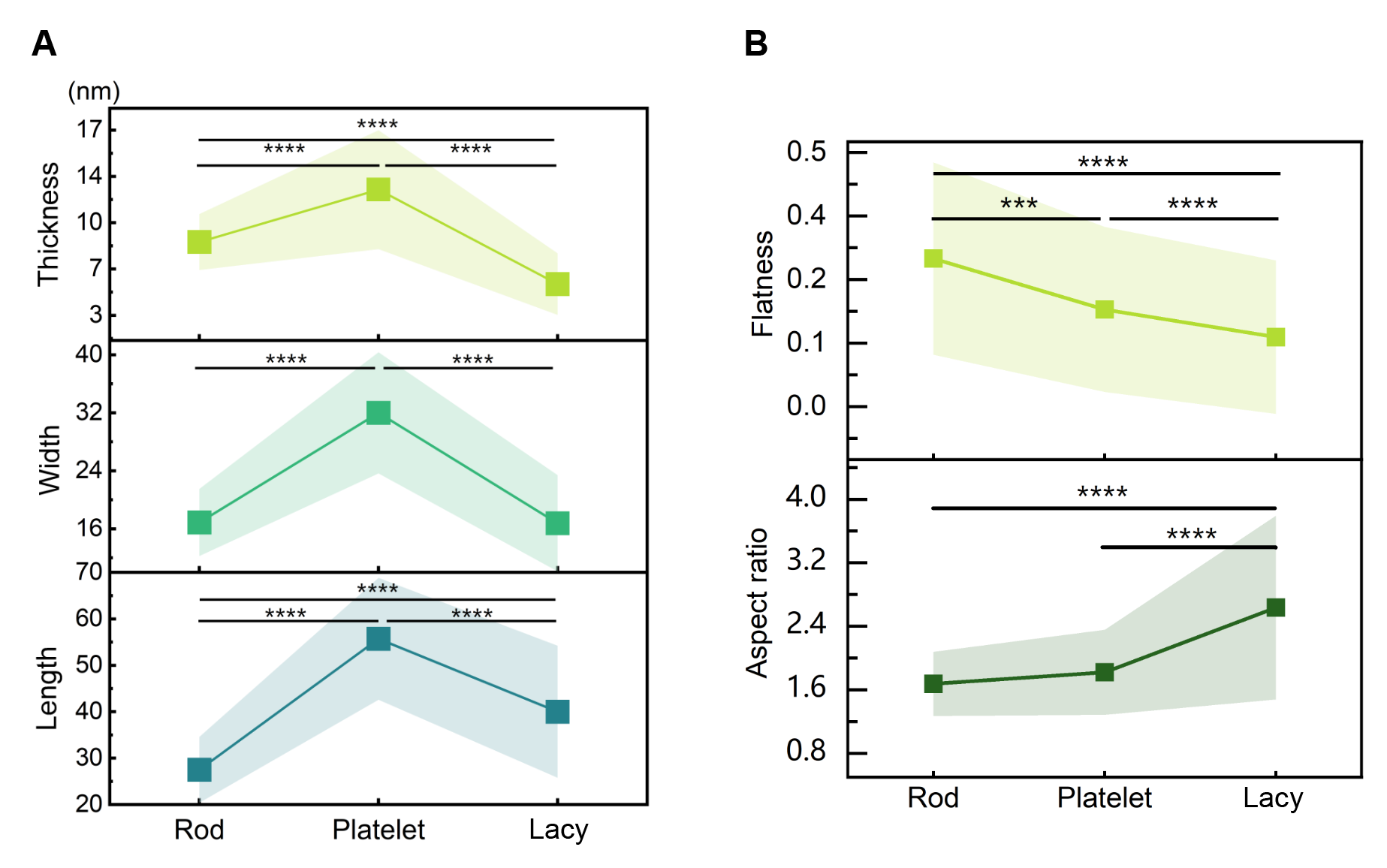

**Fig. S12.** Configurational analysis of HAP crystals in stage ii to iii based on Cryo-TEM 3D tomography. (**A**) Size of crystals shown in three dimensions. Lacy HAP crystals exhibited sizes in consistent with previously reported mature bone HAP. (**B**) Flatness and aspect ratio changes of crystals. Statistical significance is illustrated as: *p ≤ 0.05; **p ≤ 0.01; ***p ≤ 0.001.

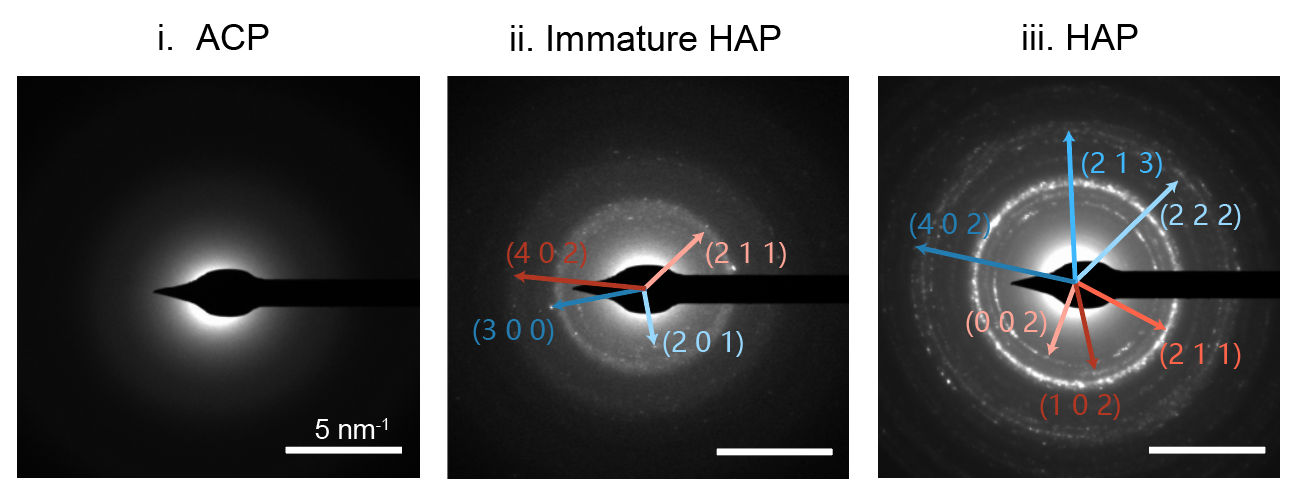

**Fig. S13.** SAED patterns of minerals in different stages. No obvious HAP lattice could be observed in ACP, while immature HAP and lacy-like HAP display the typical HAP lattice but HAP in stage iii displayed higher crystallinity. Abbreviations: SAED, selected area electron diffraction; HAP, hydroxyapatite; ACP, amorphous calcium phosphate.

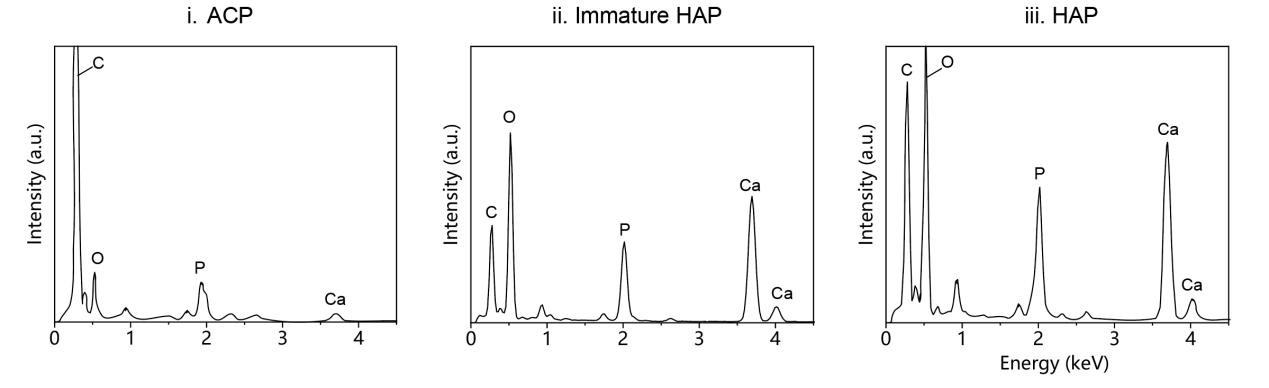

**Fig. S14.** Representative EDS spectra of mineral aggregates in three stages. Abbreviations: EDS, energy dispersive X-ray scanning.

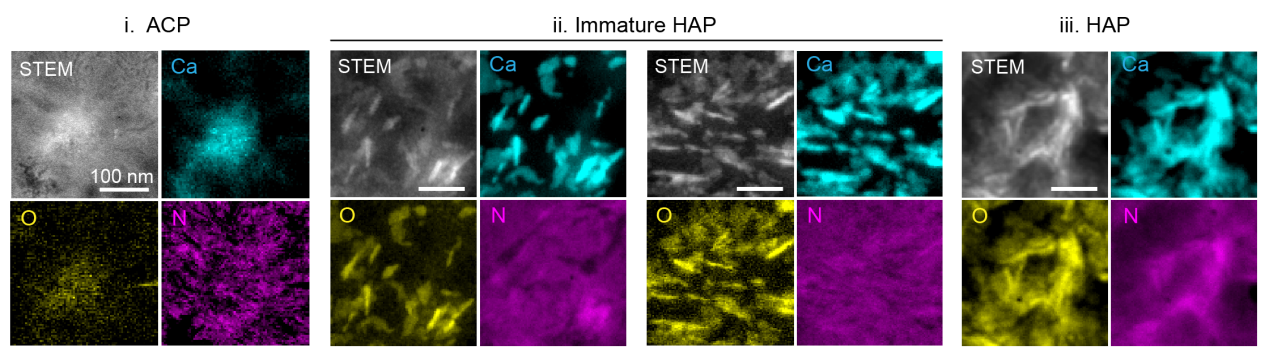

**Fig. S15.** EELS intensity maps of minerals with different configurations. STEM image and EELS intensity maps of the calcium L_2,3_ edge, oxygen K edge and nitrogen K edge are shown, corresponding to the merged images in Fig. 3h. Abbreviations: EELS, electron energy loss spectroscopy; STEM, scanning transmission electron microscopy.

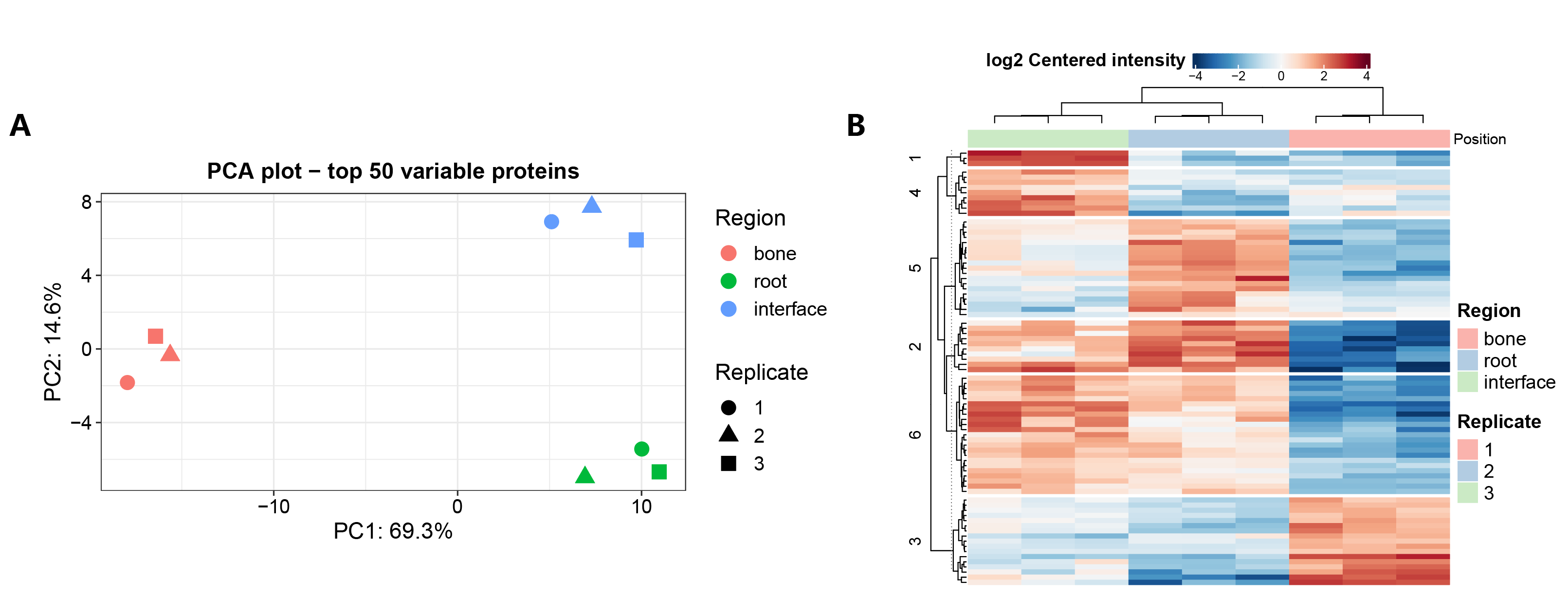

**Fig. S16.** Quantitative analysis of the human root-bone interface proteome. (**A**) Principal component analysis (PCA) clustering of root, interface, and bone tissue (n=3). (**B**) Heatmap showing differential expression of proteins in root, interface, and bone tissue. Each row represents one protein and each column represents one sample.

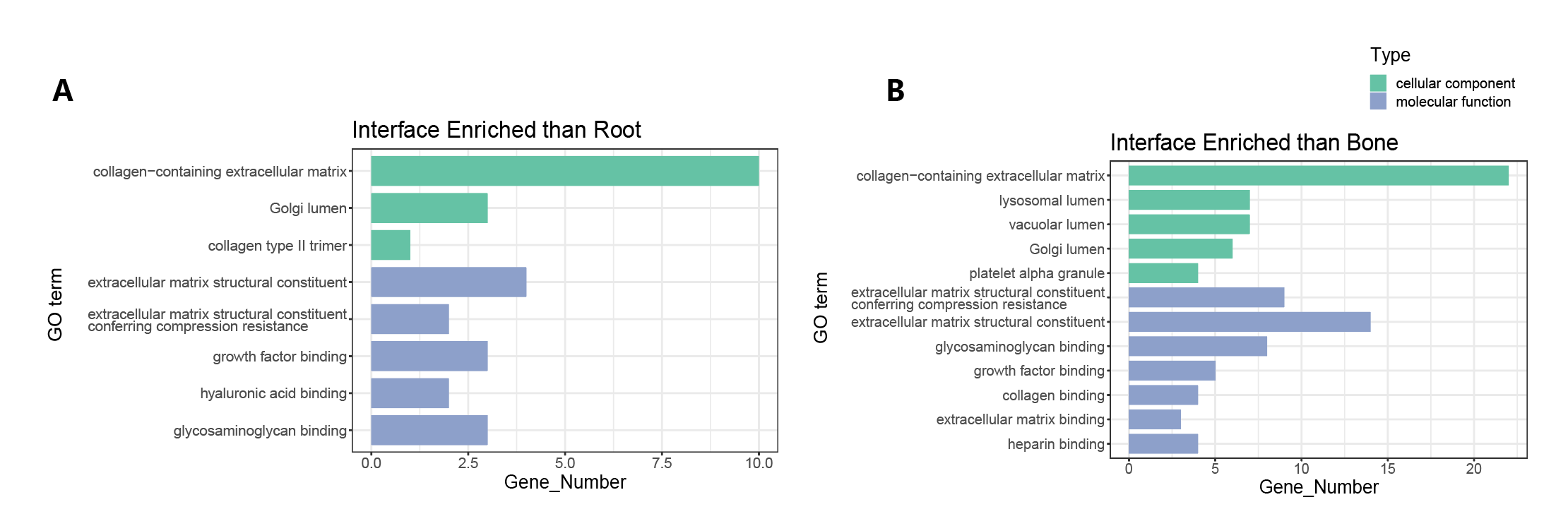

**Fig. S17.** Gene ontology (GO) enrichment analysis of proteins expressed more abundantly in the interface compared to the root and bone.

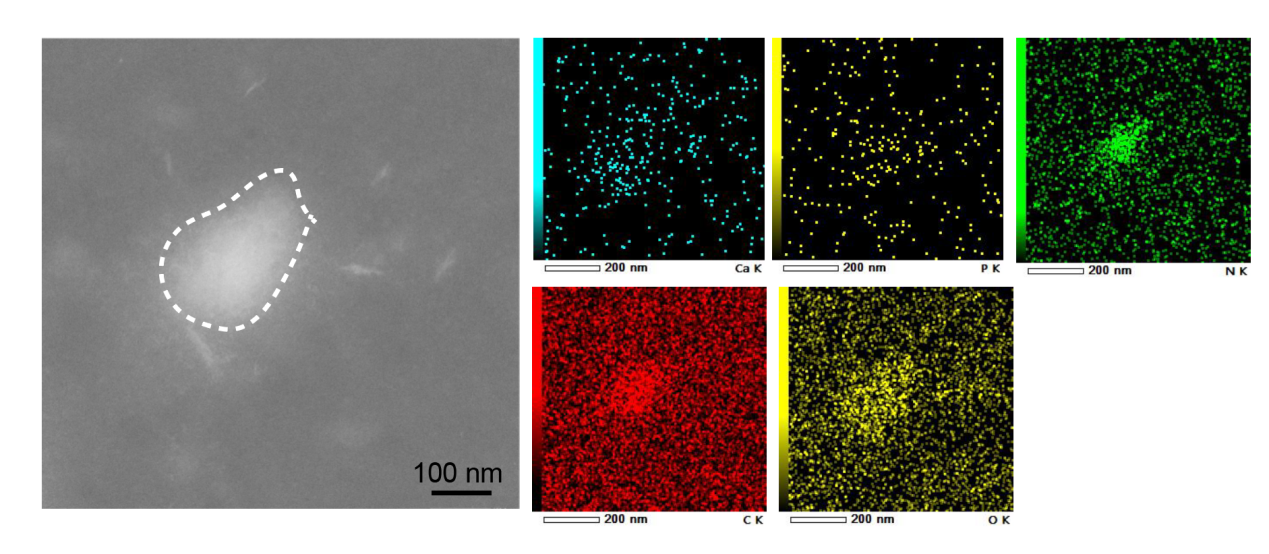

**Fig. S18.** EDS mapping of Ca, P-rich aggregate at the early stage of in vitro mineralization after treatment with MGP (day 1). The amorphous deposit outlined in a white dashed line displays a sphere-like morphology, similar to the spherical minerals found at the root-bone interface.

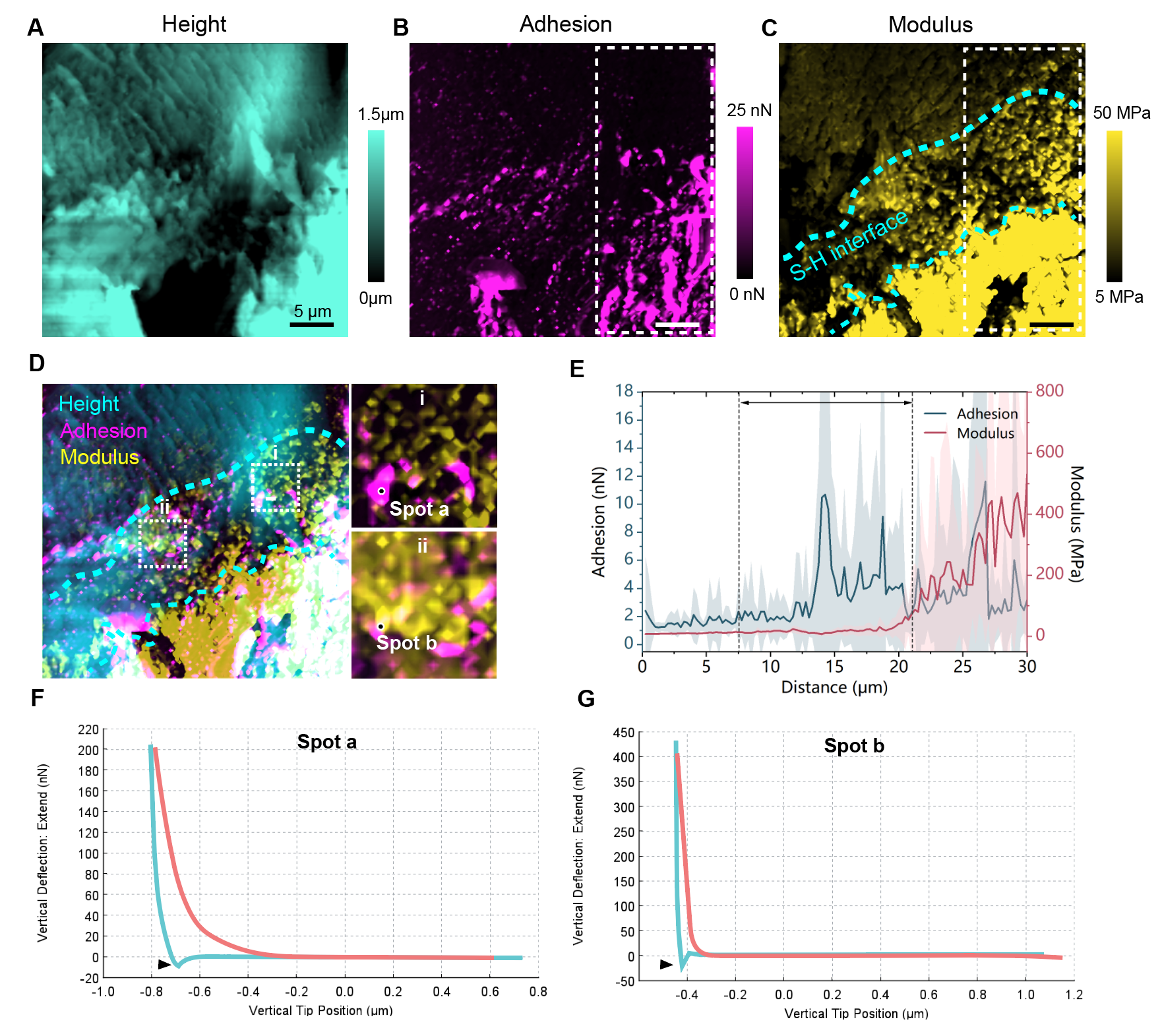

**Fig. S19.** Maps of height (**A**), adhesion (**B**), and modulus (**C**) of the S-H interface, and the merged image (**D**) showing the correlation between adhesion and tissue topography. Enlarged areas in (d) reveal an association between high-adhesion loci (spot a and b) and granules with increasing modulus. (**E**) Average modulus and adhesion force in the longitudinal direction over the S-H interface indicate that adhesion increases as modulus increases. Examples of adhesion force measured by AFM probe deflection in soft tissue (**F**) and hard tissue (**G**) are also shown.

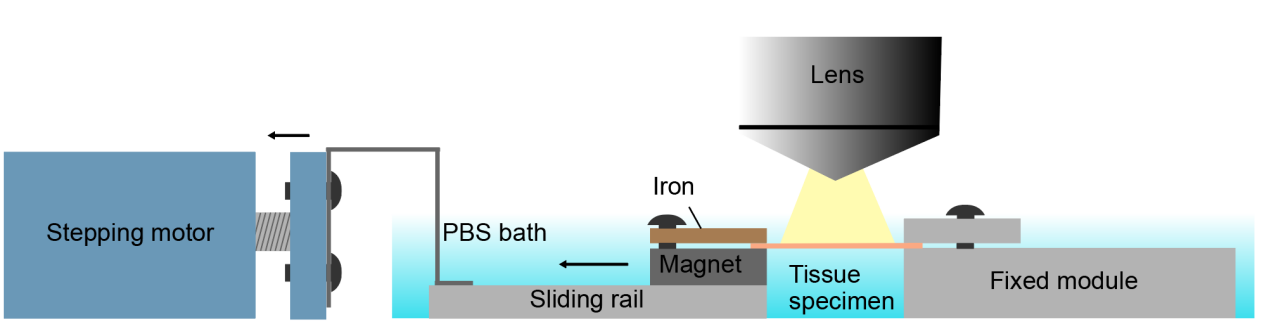

**Fig. S20.** Graphic depiction of custom-made *in situ* monitoring chamber for tissue mechanical response under fluorescent microscope.

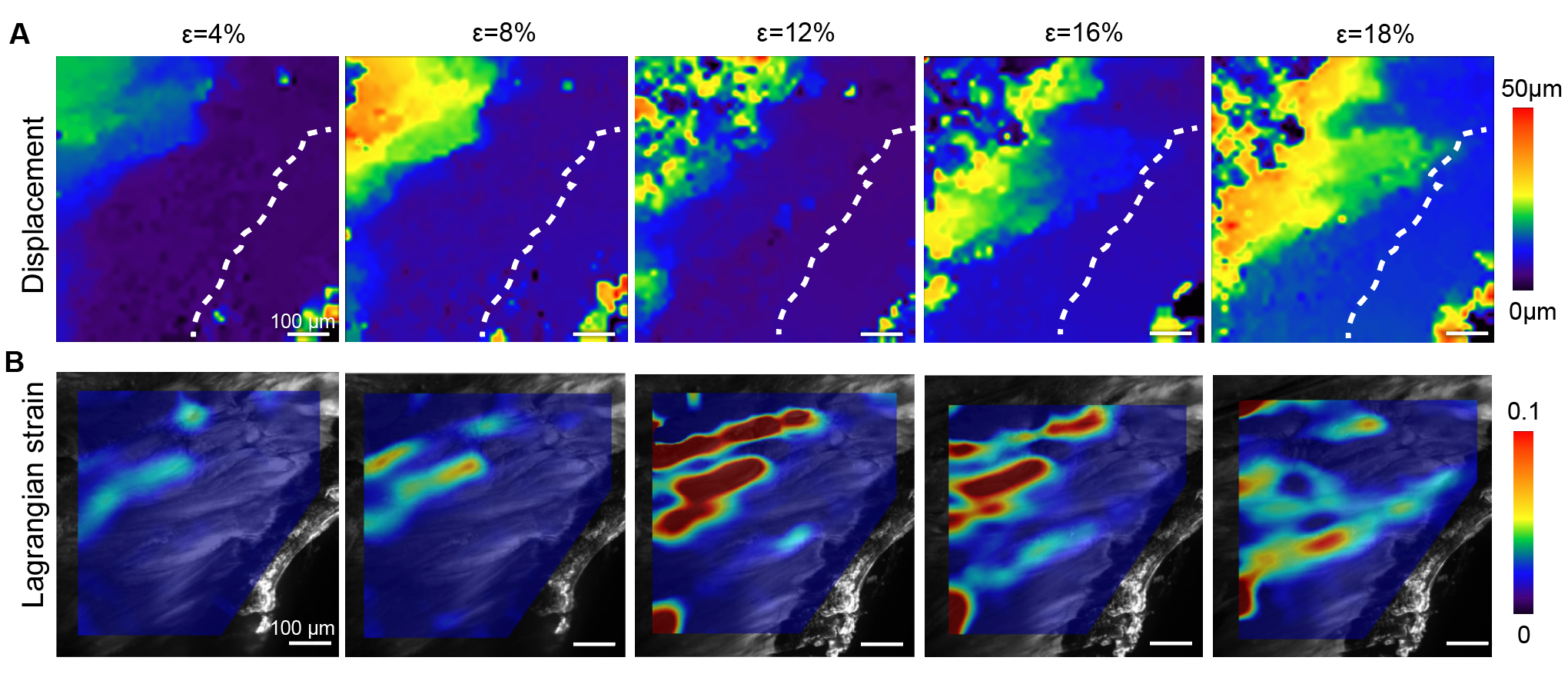

**Fig. S21.** *In situ* observation of the root-bone tissue response under tensile stress. (**A**) Displacement magnitude maps at various strain levels were obtained using the PIV algorithm. (**B**) Lagrangian strain maps at various strain levels were acquired through the DIC algorithm. Both the magnitude map and strain maps show that only the soft tissue was involved when the strain was < 15%. Abbreviations: PIV, particle iterative velocimetry; DIC, digital image correlation.

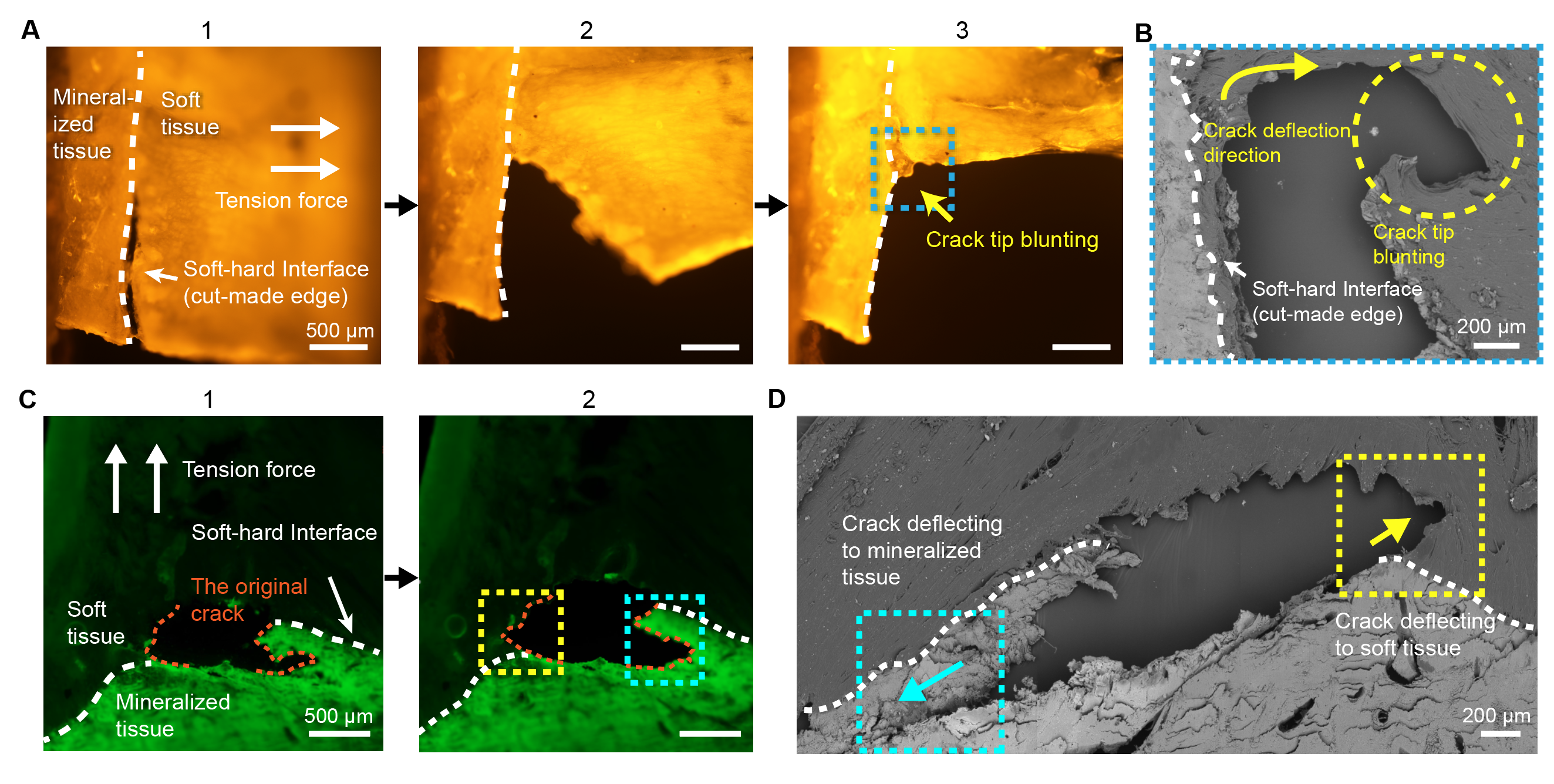

**Fig. S22.** Crack behavior on the S-H interface monitored by an in-situ monitoring chamber. (**A**) Consecutive snapshots show root-bone tissue with a pre-created crack at the edge of the S-H interface (denoted by white dashed line) deforming in response to tensile force, and the crack tip blunted as a consequence. (**B**) SEM shows the blunted crack tip (highlighted in blue dashed lines in A3) with deflection towards the soft tissue. (**C**) Consecutive photos show root-bone tissue with a pre-existing crack within the tissue at the S-H interface (denoted by orange dashed line). The crack tips in the soft tissue and mineralized tissue (outlined in yellow and blue dashed lines, respectively) displayed moderate propagation into soft and mineralized tissues rather than propagating through the interface. (**D**) SEM images revealed a similar tendency of crack propagation.

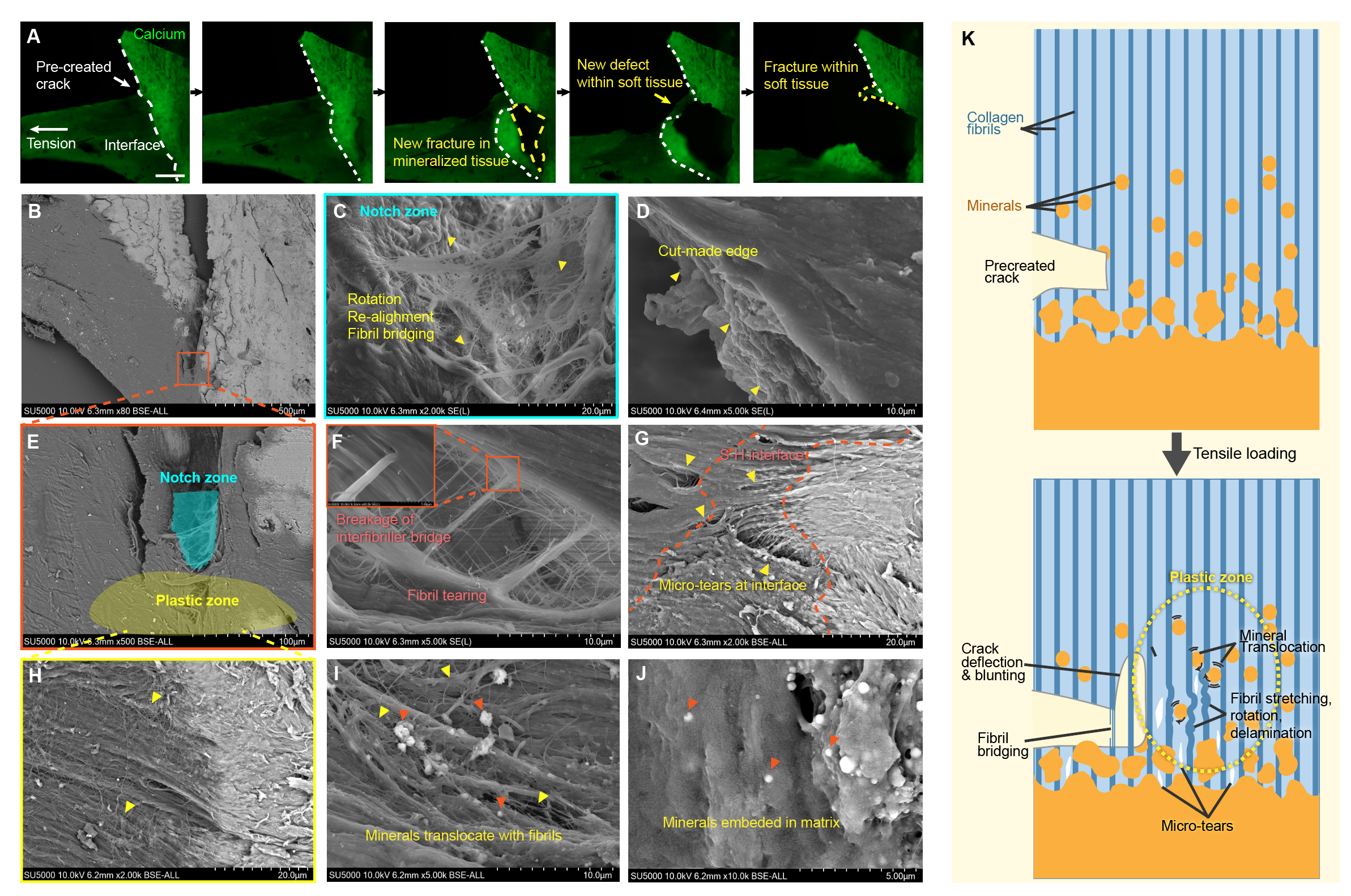

**Fig. S23.** Fracture behavior of the S-H interface. (**A**) Sequential snapshots of a tensioned root-bone tissue specimen with a pre-created crack on S-H interface. The crack propagates through the soft tissue and mineralized tissue. (**B**) SEM view of the propagated crack. The region framed in orange refers to the location of the notch and plastic zone, which are shown in (**E**). (**C**) The notch zone where fibril bridging, fibril rotation and re-alignment occurred, constituting a distinct morphology from the cut-made soft tissue edge in (**D**). (**F**) Fibril bridging region revealed fibril tearing where interfibrillar bridges broke. (G) and (H) show the S-H interface stressed in the plastic zone. The SEM image of the S-H interface within plastic zone (**G**) revealed micro-tears emerging at the unmineralized and partially mineralized region but arrested at the frontier of fully mineralized region. (**H**) Fibril re-alignment in unmineralized region around the S-H interface. (**I**) Immature mineral particles at the partially mineralized region translocate along with the re-aligned fibrils. (**J**) Immature mineral particles in non-stressed tissue were embedded in the matrix. (**K**) Graphic summary of the cooperative behavior of the S-H interface around the crack tip.

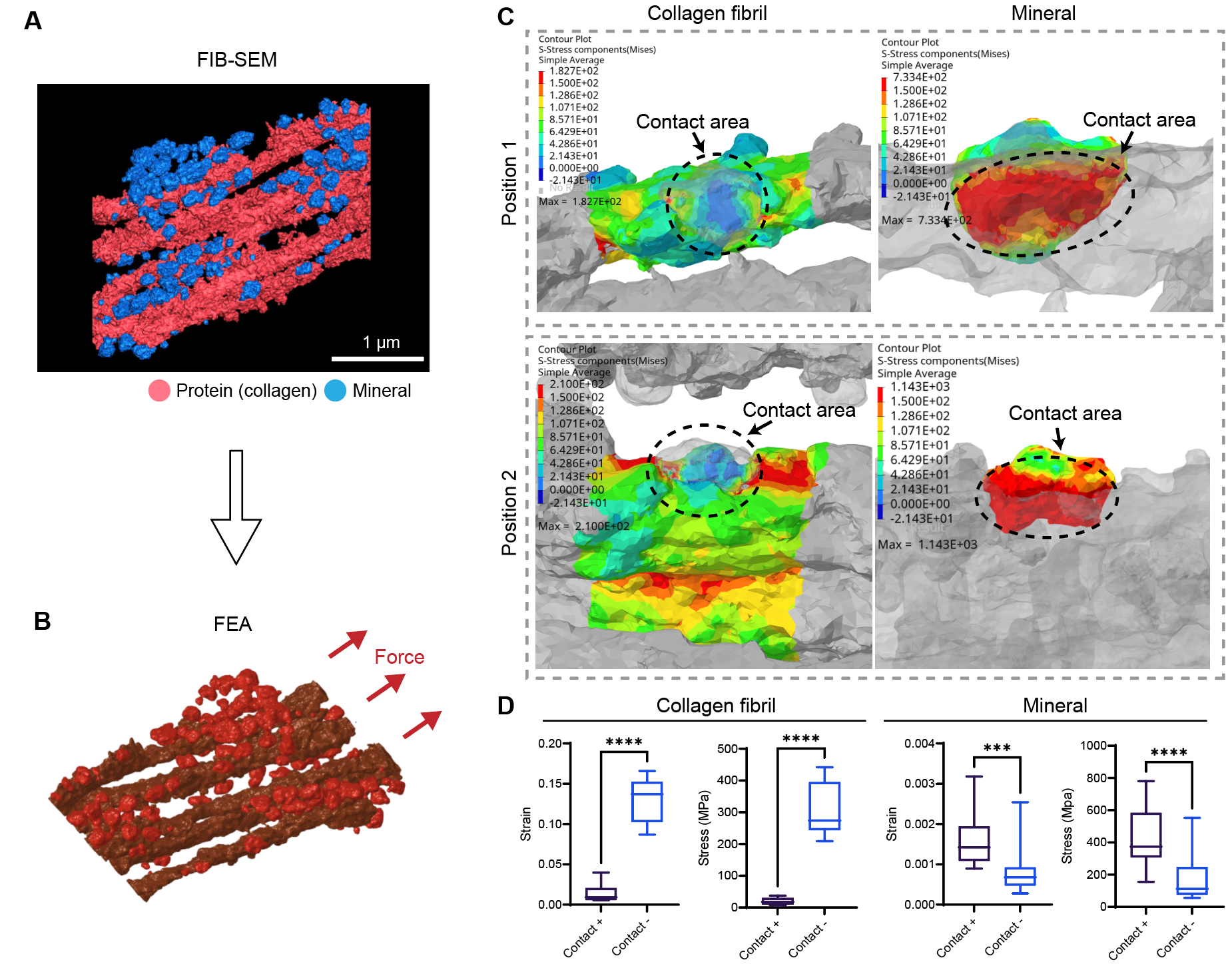

**Fig. S24.** Finite element analysis (FEA) of FIB-derived 3D microstructures of collagen fibril and immature mineral particles. (**A**-**B**) Snapshot of 3D models of collagen fibrils and mineral particles obtained from FIB-SEM and reconstructed using Amira. Minerals were located in extrafibrillar space and were in contact with fibrils. The 3D model was transferred to conduct FEA, and a tensile force in axial direction was loaded on the model. (**C**) Representative views of stress distribution on fibrils (left panel) and minerals (right panel) after tensile loading. The contact area between fibrils and minerals are indicated by black circles. (**D**) Statistical analysis of stress on contact or non-contact regions on fibrils and minerals. Statistical significance is illustrated as: *p ≤ 0.05; **p ≤ 0.01; ***p ≤ 0.001. Abbreviations: FIB-SEM, focused ion beam scanning electron microscopy.

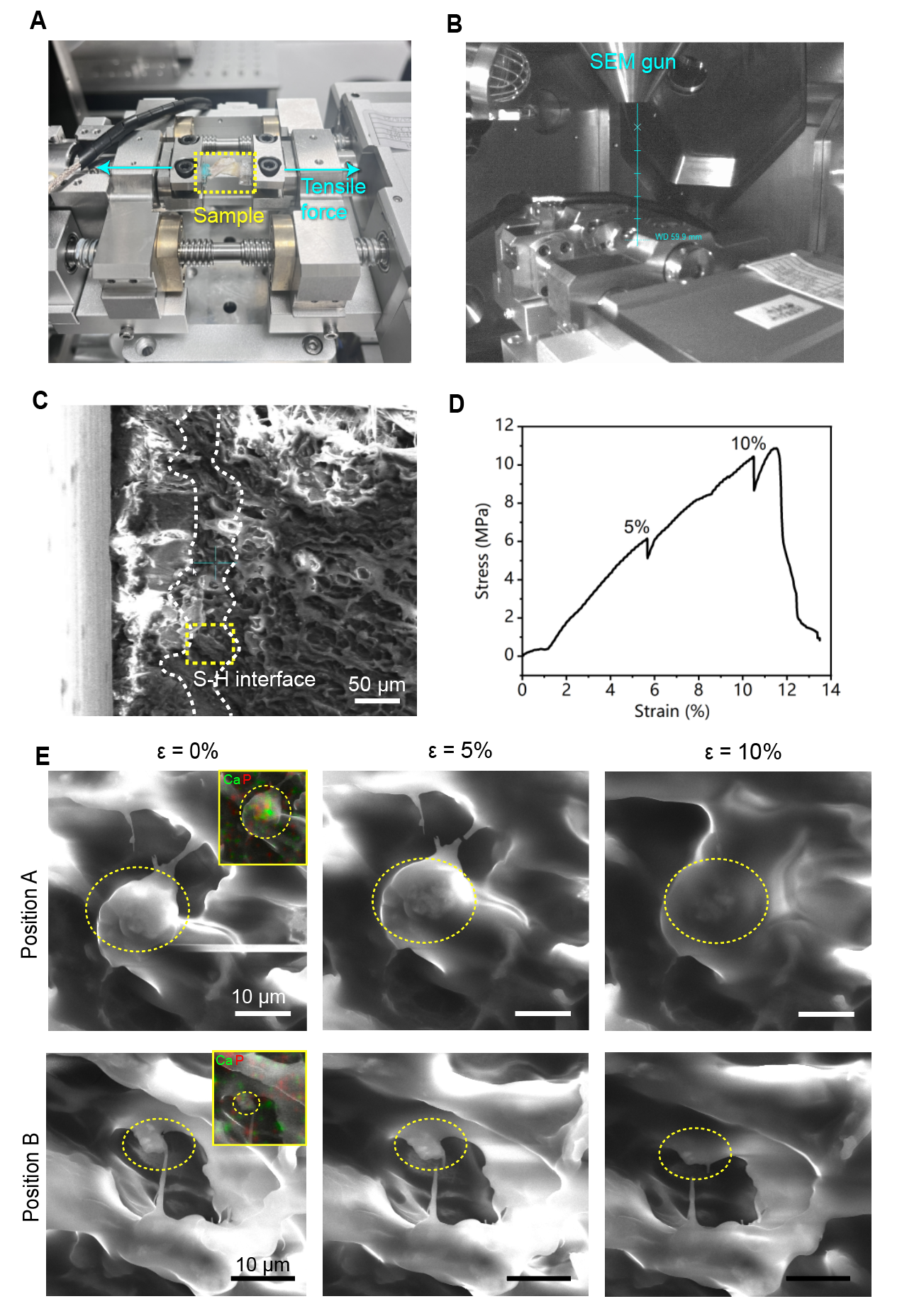

**Fig. S25.** In-situ SEM tensile test of the root-bone tissue. (**A**) Photo showing the in-situ tensile test device. (**B**) Snapshot showing the inner view of the device under the SEM gun. (**C**) SEM image of the exposed portion of the clamped root-bone tissue specimen. The S-H interface is indicated by a white dashed line. The yellow dashed line circles the monitored region-of-interest (ROI) during tensile test. (**D**) The stress-strain curve of the tested root-bone tissue. Pauses at 5% and 10% strain were for closer observation of the ROI. (**E**) The original SEM pictures of the ROI showing translocation of spherical minerals at two positions as strain increased from 0% to 5% and 10%.

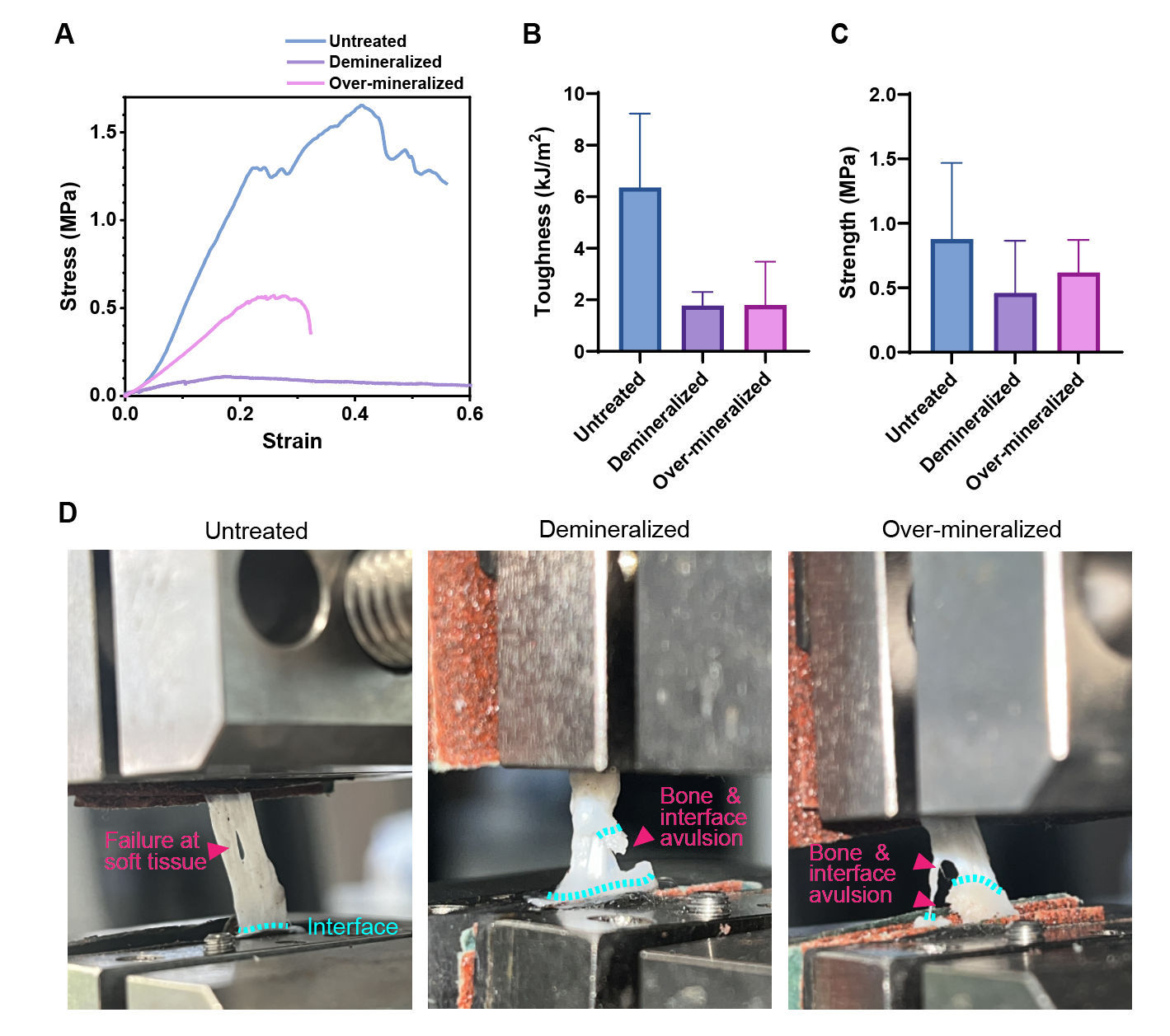

**Fig. S26**. Effect of over-mineralization and demineralization on interface mechanics. (**A**) Representative stress-strain curves of untreated, over-mineralized and demineralized root-bone samples. Demineralized samples exhibit increased ductility but diminished strength, whereas over-mineralized samples are prone to brittle fractures within the bone region. (**B**) Demineralization and over-mineralization led to tendency of reduction in interfacial toughness. (**C**) Demineralization and over-mineralization led to tendency of reduction in interfacial strength. (**D**) Failure modes after demineralization or over-mineralization were different from untreated samples. While untreated samples’ failure did not involve interface, the demineralized and over-mineralized samples were susceptible to fracture originating from bone and going through the interface.

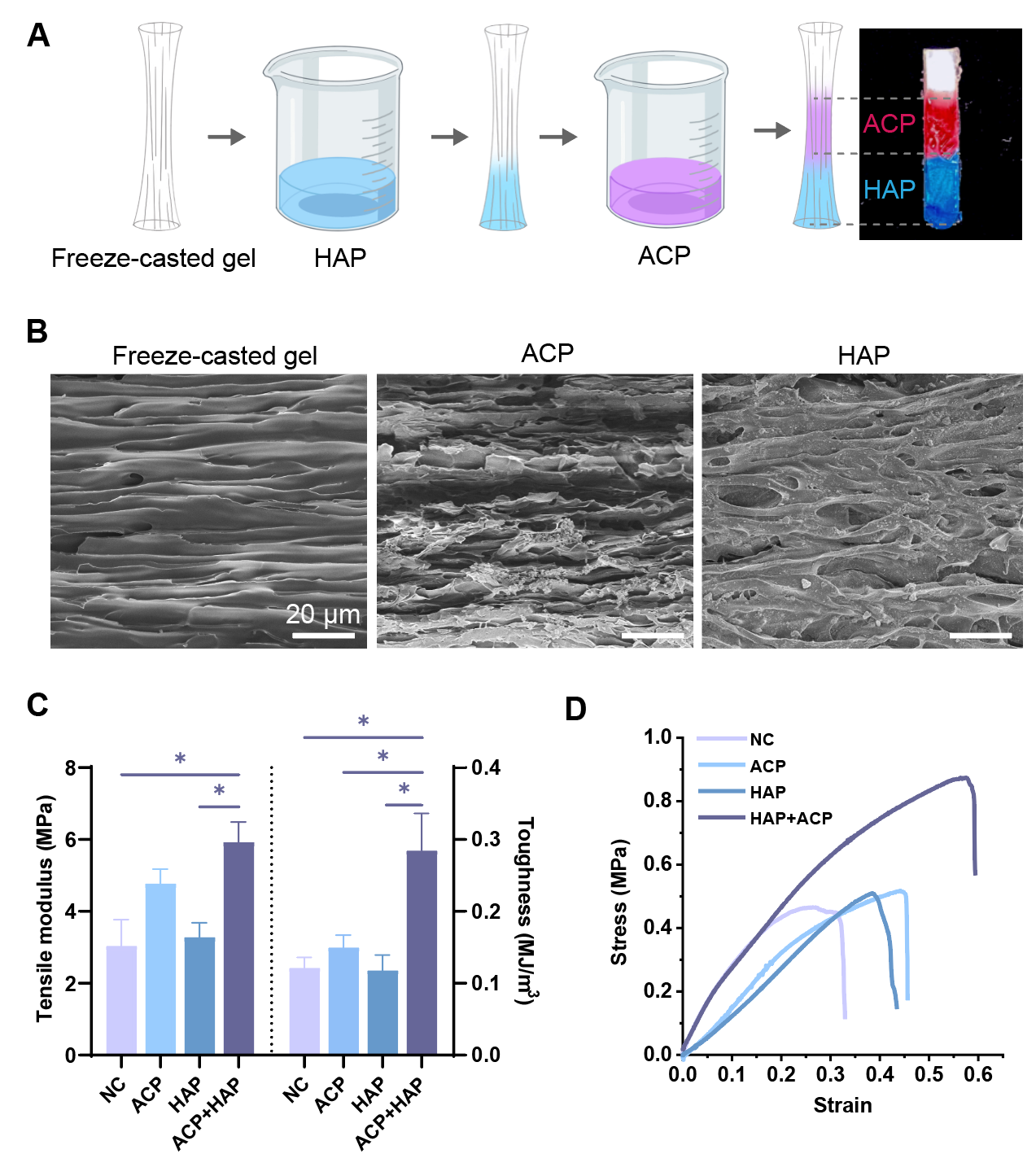

**Fig. S27.** Soft-hard interface on artificial root exhibited improved mechanical performance with ACP-HAP transformation. (**A**) Schematic of the procedure for fabricating an artificial root structure with a biomimetic transition from ACP to HAP in mineralization. (**B**) SEM snapshots of the freeze-casted aligned structure and the portion of the structure after mineralization using ACP or HAP. (**C**) Statistical analysis of the tensile modulus and toughness of four types of artificial root structures. The control group referred to artificial root that were not treated after freeze-casting or were treated with ACP or HAP only. (**D**) Representative stress-strain curves of artificial root structures in the tensile test. Abbreviations: silMA, silk fibroin methacrylate; NC, negative control. Statistical significance is illustrated as: *p ≤ 0.05; **p ≤ 0.01; ***p ≤ 0.001.

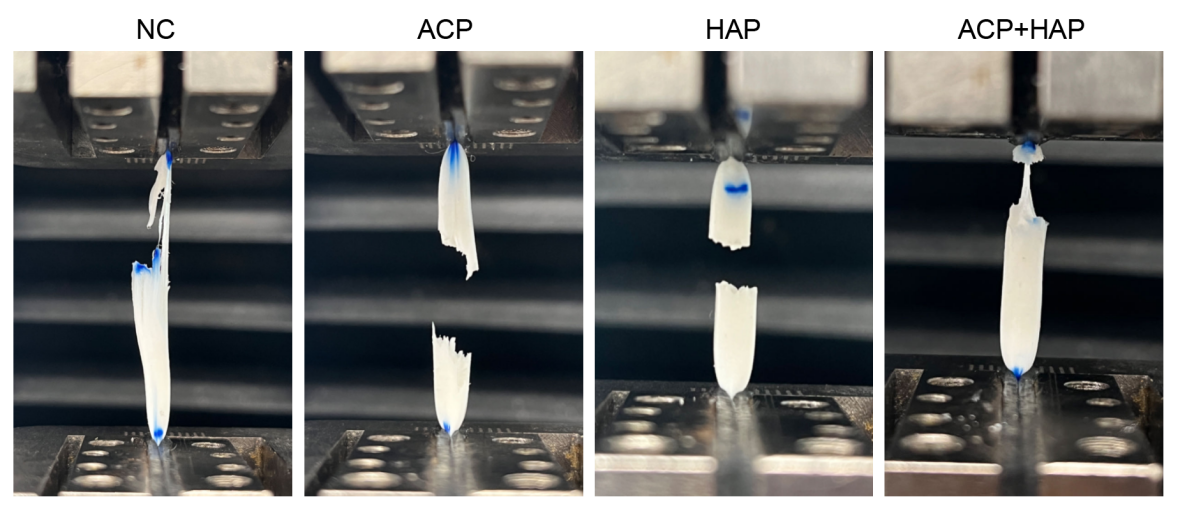

**Fig. S28.** Failure modes in the tensile test of mineralized artificial roots. Non-mineralized artificial root failed with torn fibers; ACP- and HAP-mineralized root failed at the interface between the mineralized region and the non-mineralized region; ACP+HAP artificial root failed mostly in the non-mineralized region, which had the lowest mechanical strength.

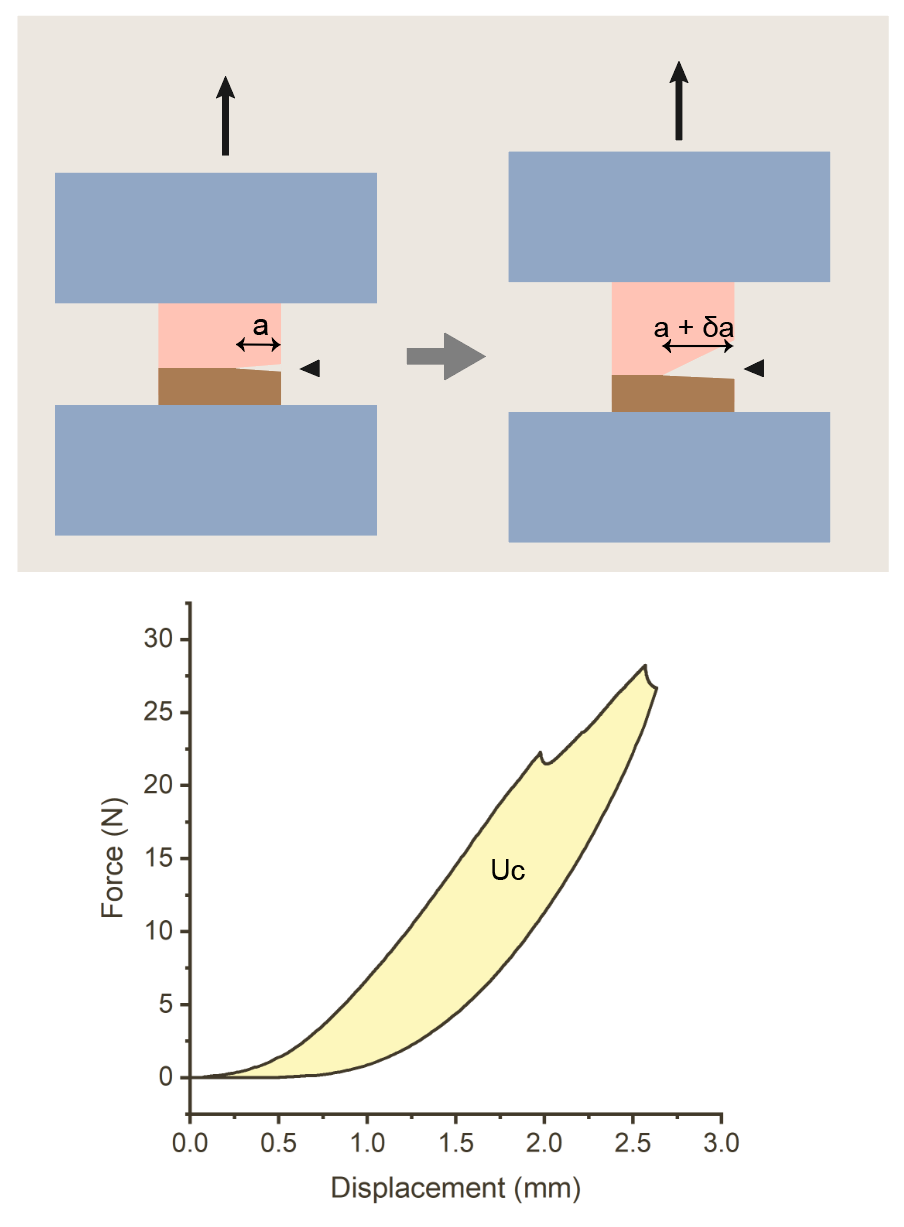

**Fig. S29.** Schematic illustration of fracture toughness estimation method.

| Donor No. | Gender | Age | Sample collected |
| --- | --- | --- | --- |
| 1 | Male | 48 | Medial & lateral, posterior & anterior roots (x4) |
| 2 | Male | 49 | Medial & lateral, posterior & anterior roots (x4) |
| 3 | Male | 39 | Medial posterior, medial anterior roots (x2) |
| 4 | Male | 37 | Medial & lateral, posterior & anterior roots (x4) |
| 5 | Male | 26 | Medial & lateral, posterior & anterior roots (x4) |

**Table S1.** Information on meniscus root-bone tissue donors.

| Stage | 1 | 2 | 3 |
| --- | --- | --- | --- |
| Mineral morphology | Spherical | Fused | Dense |
| Spatial distribution | Extrafibrillar | Extra- & partial intrafibrillar | Extra & intrafibrillar |

**Table S2.** Mineral morphological and distributional feature changes in three steps over the S-H interface.

| Raman shift (cm^-1^) | Band assignment | Component | Analyzed in | Ref. |
| --- | --- | --- | --- | --- |
| 425 | ν_2_PO_4_^3-^ | HAP | Raman | ^29^ |
| 580 | ν_4_PO_4_^3-^ | HAP | Raman | ^29^ |
| 952 | PO_4_^3-^ (ACP) | ACP | Raman &SRS | ^60^ |
| 960 | ν_1_PO_4_^3-^ (HAP) | HAP | Raman &SRS | ^61^ |
| 1065-1075 | ν_1_CO_3_^2-^ | Carbonate group | Raman | ^61^ |
| 1410 | νs(COO^-^) | GAG | SRS | ^59^ |
| 1440 | δ(CH_2_) | Lipid | Raman &SRS | ^62^ |
| 1450 | δ(CH_2_) | Collagen (proteins) | SRS | ^59^ |
| 1668 | Amide I | Collagen (proteins) | Raman | ^59^ |
| 2850 | CH_2_ stretch | Lipids | SRS | ^63^ |
| 2926 | CH_3_ stretch | Proteins | SRS | ^63^ |

**Table S3.** Raman spectroscopy and SRS band assignments for the analysis of the S-H interface.

| **Elements** | **Approximate energy loss (eV)** | **Peak description** |
| --- | --- | --- |
| Phosphorus | | |
| a | 138 | Transitions to p-like states |
| b | 141 | Transitions from 2p state to new state from interactions with calcium 3D orbital function (characteristic for calcium-containing minerals) |
| c | 146 | Transitions to d-like states |
| d | 160 | Multiple scattering and the maximum 2p state cross section |
| Carbon | | |
| e | 284.2 | 1s to π* (C=C)/ Amorphous carbon probably from the embedding resin |
| f | 287 | Transitions to the vacant π* state if carbonyl groups probably from amino acids |
| g | 290 | 1s to π* (C=O)/CO3 groups |
| h | 296.7 | 1s to σ* (C-C)/ Amorphous carbon |
| Calcium | | |
| i | 348 | Transitions from L3(2P3/2) to d-like states |
| j | 351 | Transitions from L2(2p1/2) to d-like states |
| Nitrogen | | |
| k | 400-401 | 1s-π* transitions (characteristic of collagen crosslinking) |
| l | 408 | 1s-σ* transitions in amino compounds |
| Oxygen | | |
| m | 537 | Transitions to the vacant π* states of the Ca-O bonding environment |
| n | 539 | Transitions to the vacant σ* states of the Ca-O bonding environment |
| o | 545 | Transitions to 4s- and 4p-like states in calcium-oxygen bonds |

**Table S4.** EELS characteristic peaks of minerals, organics, and resin at the S-H interface.

| Iteration | Interrogation window size (px) | Search window size (px) | Vector spacing (px) |
| --- | --- | --- | --- |
| 1 | 128 | 64 | 48 |
| 2 | 256 | 128 | 128 |
| 3 | 64 | 32 | 16 |

**Table S5.** Parameters used in the PIV algorithm for the analysis of displacement magnitude.

**Movie S1. (separate file)**

The FEA result of a S-H interface two-dimensional model being tensioned.

**Movie S2. (separate file)**

3D reconstruction of a collagen fibril-mineral complex in interfibrillar mineralization area. Data originates from FIB-SEM images.

**Movie S3. (separate file)**

Animated video highlighting the toughening mechanisms of the human meniscus root-bone interface.

**Data S1. (separate file)**

Full results of proteomic analysis of root-bone interface. Proteins enriched at interface region were categorized according to function summary on Genecards.org.

**Supplementary information references**
